# Cross-frequency slow oscillation–spindle coupling in a biophysically realistic thalamocortical neural mass model

**DOI:** 10.1101/2021.08.29.458101

**Authors:** Nikola Jajcay, Caglar Cakan, Klaus Obermayer

## Abstract

Sleep manifests itself by the spontaneous emergence of characteristic oscillatory rhythms, which often timelock and are implicated in the memory formation. Here, we analyze a neural mass model of the thalamocortical loop of which the cortical node can generate slow oscillations (approx. 1 Hz) while its thalamic component can generate fast sleep spindles of *σ*-band activity (12–15 Hz). We study the dynamics for different coupling strengths between the thalamic and cortical nodes, for different conductance values of the thalamic node’s potassium leak and anomalous rectifying currents, and for different parameter regimes of the cortical node. The latter are: (1) a low activity (DOWN) state with noise-induced, transient excursions into a high activity (UP) state, (2) an adaptation induced slow oscillation limit cycle with alternating UP and DOWN states, and (3) a high activity (UP) state with noise-induced, transient excursions into the low activity (DOWN) state. During UP states, thalamic spindling is abolished or reduced. During DOWN states, the thalamic node generates sleep spindles, which in turn can cause DOWN to UP transitions in the cortical node. Consequently, this leads to spindle-induced UP state transitions in parameter regime (1), thalamic spindles induced in some but not all DOWN states in regime (2), and thalamic spindles following UP to DOWN transitions in regime (3). The spindle-induced *σ*-band activity in the cortical node, however, is typically strongest during the UP state, which follows a DOWN state “window of opportunity” for spindling. When the cortical node is parametrized in regime (3), the model well explains the interactions between slow oscillations and sleep spindles observed experimentally during Non-Rapid Eye Movement sleep. The model is computationally efficient and can be integrated into large-scale modeling frameworks to study spatial aspects like sleep wave propagation.

## 1. Introduction

Sleep marks a pronounced change of the brain state as one of the vital means of persisting mental and physical health [1]. It manifests itself by the spontaneous emergence of characteristic oscillatory rhythms, most visible in the electroencephalogram (EEG) but also noticeable in intracellular recordings, electrooculography (EOG), and electromyography (EMG) [2, 3]. Distinct oscillatory features form the basis for sleep classification into several stages: rapid eye movement sleep (REM) and three stages of non-REM sleep (N1 through N3) [4, 5]. The NREM sleep stages exhibit characteristic patterns: the N1 stage consists of slow eye movements and low-amplitude low-frequency *δ*-band EEG activity [6]; sleep spindle oscillations dominate the N2 stage with a waxing and waning envelope and an underlying oscillation in the *σ*-band (~12–15 Hz) [7], in the case of “fast” spindles, or ~9–12 Hz for “slow” spindles [8, 9]. Sleep spindles are also observed in the deeper, N3 sleep stage, albeit with lower power in the fast spindle frequency range [10] and lower density [11]. Finally, slow oscillations (SOs), which is an alteration of active (UP) and silent (DOWN) cortical states at ~1 Hz frequency, governs the deepest N3 sleep stage but is also present in the N1 and N2 sleep stages, albeit with a lower spectral power [12, 13]. Ripple oscillations are the second hallmark of the N3 sleep stage. Ripples are fast oscillations (80–140 Hz) that occur in hippocampal networks, often accompanied by a sharp wave, and signify reactivations (memory replay) of neural ensembles in these networks [14, 15]. The precise coordination of the slow oscillations, spindles, and ripples was shown to be vital for memory consolidation, of which the main manifestation is the reactivation of specific activity patterns — i.e., memory replay — during sleep [16, 17, 18, 19]. The hierarchical nesting of spindle waxing periods to the depolarized cortical UP states, mediated by the thalamocortical circuitry, is essential for consolidation by providing a “window of opportunity” and favorable conditions for plasticity for transferring episodic memories from short-term hippocampal storage to longer-term neocortical storage [20, 9, 21]. Notably, it was recently shown that the interplay of the aforementioned rhythms is vital also for non-hippocampus-dependent consolidation [15] in rodents [22] and humans [23].

The basis of slow oscillations consists of a widespread alternation of hyperpolarization and depolarization activity in neocortical networks [24, 7, 25]. In contrast, spindles are generated by the interaction of inhibitory reticular thalamic and excitatory thalamocortical neurons [26]. Spindles occur in the isolated thalamus both *in vivo* and *in vitro* [27, 28]. However, the cortex can also become actively involved in their initiation and termination [29], as well as their long-range synchronization [30, 31]. Therefore, in the thalamocortical modeling, the assumption is that the cortical part of the model generates slow oscillations and the thalamic part of the model, in turn, generates spindles [32, 33, 34]. Moreover, slow oscillations induce thalamic spindles, which then reflect back to the cortex [35]. Furthermore, it has been shown that the phase of SOs modulated the spindle power such that it exhibited UP and DOWN states similar to SOs themselves, with positive peaks during the depolarizing SO UP state, close to (or slightly before) the SO peak [9, 36].

Due to the computational advantage, the ability to elucidate the dynamical repertoire, and the ability to describe macroscopic phenomena comparable with neuroimaging datasets [37, 38], we focus here on neural mass models (NMMs hereafter). In particular, Suffczynski et al. [33] proposed a thalamocortical NMM to explain the relationship between spindle-generating and spike-wave-generating activity. More recently, Cona et al. [39] proposed a new NMM to describe a “sleeping” thalamocortical system with tonic and bursting firing modes within thalamic neurons, while Costa et al. [34] probed a thalamocortical model that generates spindles and K-complexes. To this date, however, no study focused on the cross-frequency slow oscillation–spindle interaction in the model setting that would mimic the empirical findings [40, 41, 42]. We aim to close this gap with the present study.

This study effectively extends the results in [34] by using a biophysically realistic model for the cortical node capable of generating *in vivo*-like slow oscillations [43]. By merging two modeling approaches, we also highlight the applicability of hybrid modeling approaches and their advantages, mainly the extensibility with other mass models. In our case, using an already explored cortical node facilitates the exploration of biophysically realistic stimulation protocols of the thalamocortical model. We present a thorough dynamical investigation of the thalamocortical model in the NREM sleep setting. Both cortical and thalamic modules are biophysically realistic while adhering to the notion of mass models and keeping their computational advantages. In addition to investigating the dynamical landscape, we study the nature of cross-frequency coupling between slow oscillations and sleep spindles, which are believed to set the stage for successful memory consolidation and transfer from the hippocampus to the neocortex [44, 45].

The following sections first introduce the thalamic and cortical models, their basic dynamical properties, and their respective state spaces. Next, we investigate the effects of perturbations on both isolated models to understand their response to external stimuli. Finally, we study the full thalamocortical loop, including both feedforward and feedback connections. In the full model investigation, we focus on the interaction between cortical slow oscillations and thalamic spindles, the cause-effect mechanical understanding in the spindle and SO generation and timing, and, finally, their cross-frequency coupling.

## 2. Materials and methods

### 2.1. Model design

The architecture of the thalamocortical model is shown in Fig. 1. It consists of two thalamic neural populations, namely thalamocortical neurons (TCR) and thalamic reticular nucleus (TRN), which act as excitatory and inhibitory populations, respectively. The cortical module consists of mean-field approximation of excitatory and inhibitory exponential integrate-and-fire neurons grouped into two populations, of which the excitatory subpopulation exhibits a somatic spike-frequency adaptation mechanism. Both thalamic and cortical nodes rely on the notion of an empirical firing rate which effectively replaces the complex individual spiking dynamics of neural populations.

**Figure 1:**
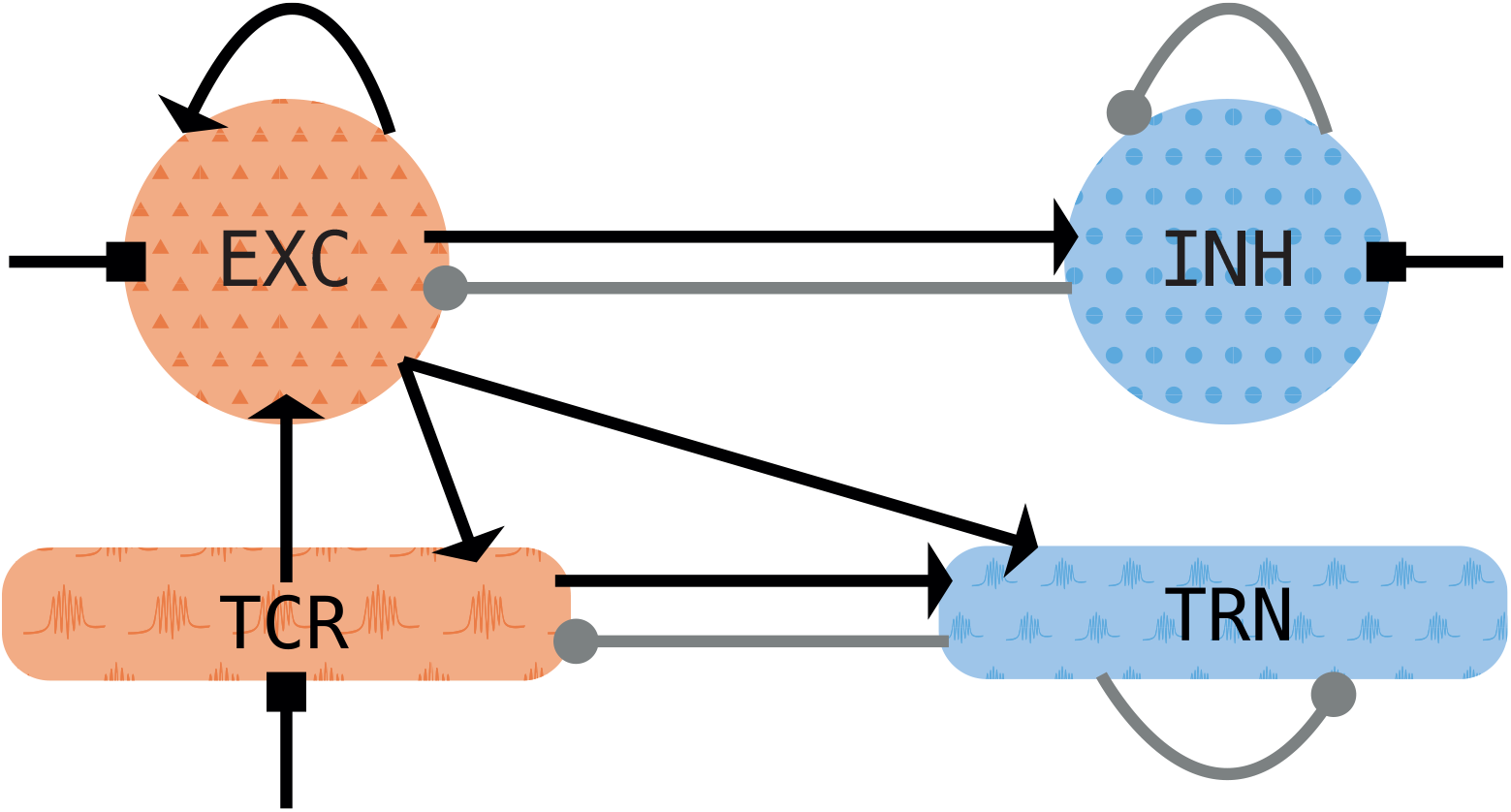
Schematic of the thalamocortical motif. The cortical node (top row) consists of one excitatory (*EXC*) and one inhibitory (*INH*) population. It is coupled to the thalamic node with its thalamocortical relay (*TCR*) and thalamic reticular nuclei (*TRN*) populations. Excitatory (inhibitory) populations are shown in orange (blue). Excitatory synapses are depicted with arrows, inhibitory synapses with filled circles. Squares depict noisy background inputs.

The model connectivity, both feedback and feedforward, relies on the fast ionotropic excitatory and inhibitory synapses conveyed by the AMPA and GABA_A_ receptors. The cortical node has feedback and feedforward connections, while in the thalamic module, only the TRN possesses feedback connections (see Fig. 1) since thalamic relay cells generally do not form local connections within the population [46].

For the connection between the thalamus and the cortex, we assume that the long-range afferents from the cortical excitatory population project to both thalamic populations and that the TCR population projects to the cortical excitatory population, as depicted in Fig. 1. We set delays of these long-range connections to a physiologically realistic value of 13 ms [47, 34].

### 2.2. Thalamic model

The thalamic model follows the approach detailed in Costa et al. [34]. The evolution of the mean membrane potential *V_α_* of the thalamic populations *α* ∈ {*t, r*} is described by

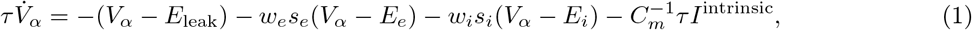

with membrane time constant *τ*, synaptic input rate *w_e_* (*w_i_*) that scales synaptic inputs *s_e_* (*s_i_*) for excitatory (*e*) and inhibitory (*i*) synapses, the corresponding Nernst reversal potential *E_e_* (*E_i_*), and the membrane capacitance *C_m_*. The mean membrane potential *V_α_* is then converted to a firing rate *r*(*V*) by a sigmoidal transfer function,

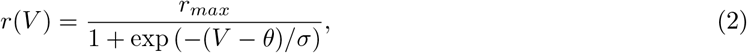

with maximum firing rate *r_max_*, firing threshold *θ*, and gain coefficient *σ*.

Both thalamic populations contain additional intrinsic currents (*I*^intrinsic^ in Eq. (1)), because spindle oscillations require rebound burst activity. Rebound bursting is impossible with a monotonic firing rate function and demands additional mechanisms. Following [34], we employ the Hodgkin-Huxley-type extension, which has been derived from integrate-and-fire-or-burst neurons [48].

The intrinsic currents in both thalamic populations include a potassium leak current,

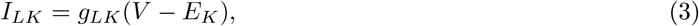

and a T-type calcium current,

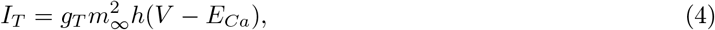

which de-inactivates upon depolarization. In both definitions, *g_LK_* and *g_T_* denote the conductance of the respective intrinsic current, *E_K_* and *E_Ca_* their respective Nernst reversal potential, and 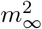 and *h* the gating functions of T-type calcium current. Both currents are essential for the generation of low-threshold spikes and rebound bursts (see Fig. 2). The definition of *I_T_* for TRN follows [49], while *I_T_* within the TCR population is given in [50]. In addition, the TCR population contains a hyperpolarization-activated cation-nonselective current *I_h_*:

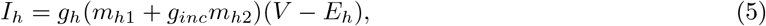

with *g_h_* being its conductance, *E_h_* its Nernst reversal potential, *m*_*h*1_ and *m*_*h*2_ the gating functions, and *g_inc_* the conductivity scaling. This hyperpolarization-activated current is responsible for the waxing and waning of spindle oscillations in the isolated thalamus [51].

**Figure 2:**
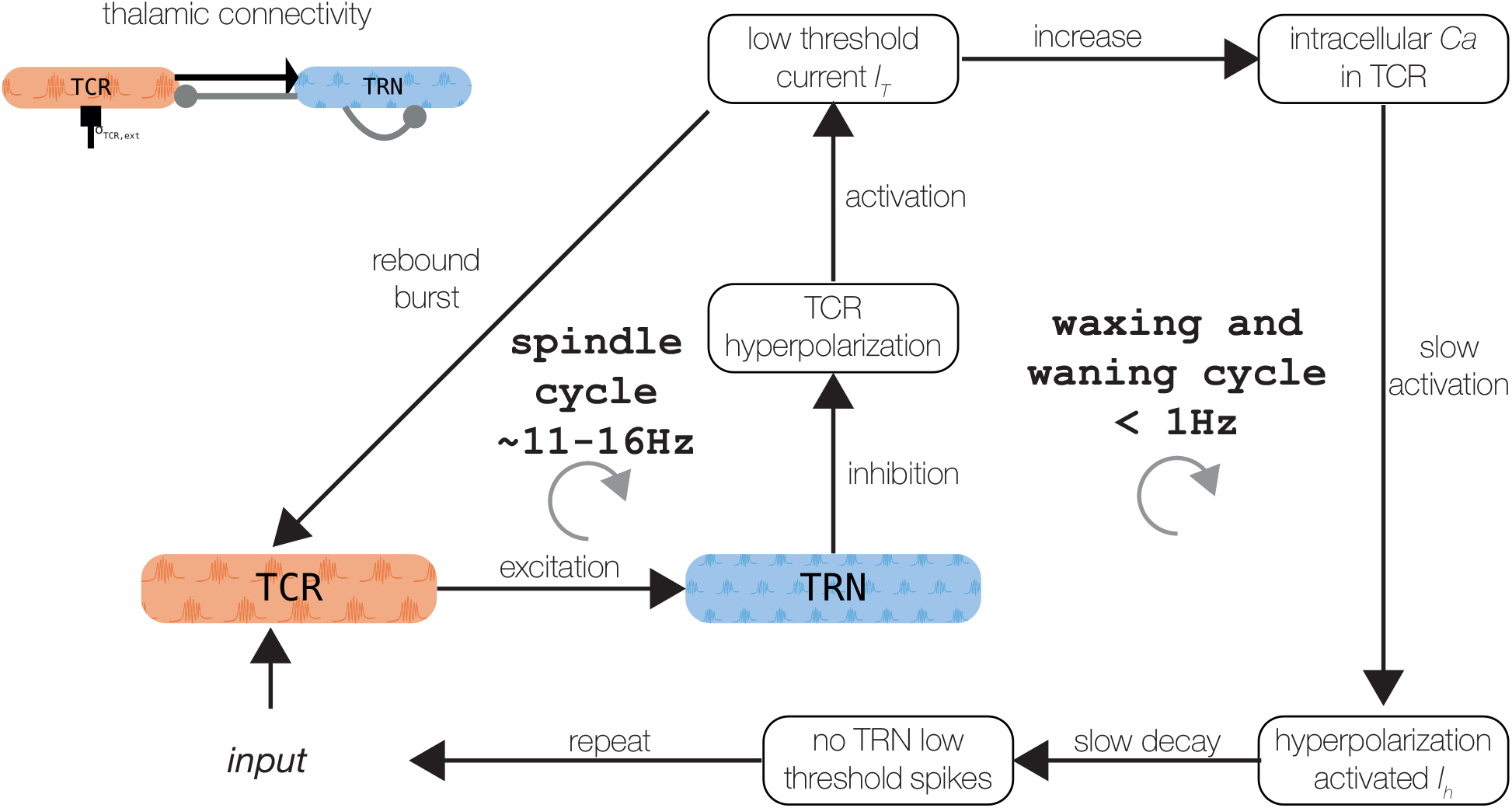
Schematic of the spindle generation mechanisms. See the text for a detailed explanation. Thalamic connectivity is shown in the upper left panel. The figure was adapted from Mayer et al. [74].

Synaptic transmission in the thalamic model is conveyed by conductance-based synapses, where the spike rate *r_k′_* of a presynaptic population *k′* elicits a postsynaptic response *s_lk_* in population *k* given by

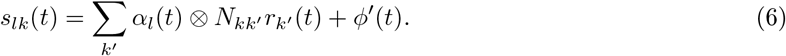

*N_kk′_* is the connection strength between the presynaptic, *k′*, and postsynaptic, *k*, populations. *ϕ′*(*t*) represents a background noise input, ⊗ denotes a convolution, and *α_l_*(*t*) is an alpha function given by

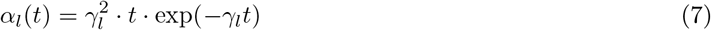

representing the synaptic response to a single spike. *γ_l_* is the decay constant of the synaptic response, and *l* ∈ {*e, i*} denotes the type of synapse, i.e., excitatory AMPA or inhibitory GABA_A_. The convolution in Eq. (6) is replaced in the numerical simulations by the second-order ordinary differential equation (ODE)

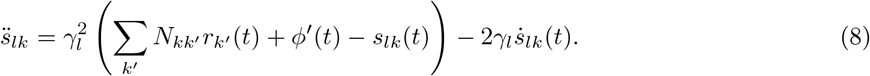

The background noise input *ϕ′*(*t*) is modeled as an Ornstein-Uhlenback process with zero drift but finite variance, and represents unresolved processes in our model, e.g. afferents from other brain regions that are not explicitly modeled here.

In summary, the thalamic node is described by the set of equations

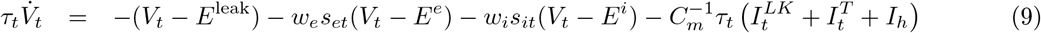

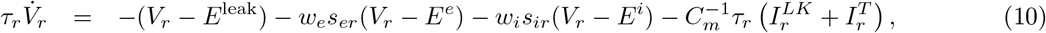

where subscripts *t* (*r*) represent the TCR (TRN) population. Note that the background noise is included on the excitatory synaptic input at the TCR population, *s_et_*. The full set of equations is given in Appendix F, with all parameters summarized in Table G.1.

### 2.3. Cortical model

The adaptive exponential integrate-and-fire (AdEx) neuron model [52] forms the basis for the derivation of the cortical mass model. Each population *α* ∈ {*E, I*} possesses *N_α_* neurons and the membrane voltage of neuron *j* in the population *α* is governed by

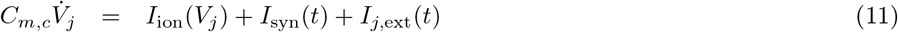

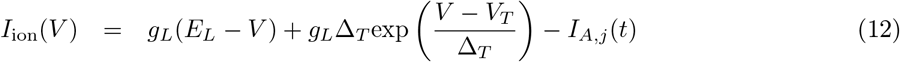

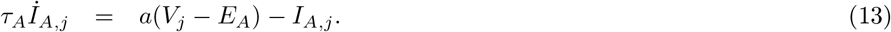

The first equation describes the temporal evolution of neuron’s *j* membrane voltage *V_j_* as a function of its internal current dynamics conveyed by *I*_ion_(*V_j_*), its synaptic current *I*_syn_(*t*), and a background current received from neural populations not specificed by the computational model external current *I*_*j*,ext_(*t*). *C_m,c_* denotes the cortical neuron’s membrane capacitance. Note, that inhibitory population *α* = *I* does not include the adaptation current. The first term in Eq. (12) expresses the voltage-dependent leak current with leak conductance *g_L_* and leak reversal potential *E_L_*, the second term describes the exponential spike initiation mechanism with slope factor Δ_*T*_ and exponential threshold *V_T_*. Finally, the last term describes the somatic adaptation current, *I_A,j_*(*t*) (Eq. 13), with subthreshold adaptation *a*, adaptation reversal potential *E_A_*, and adaptation time scale *τ_A_*. When the membrane voltage crosses the spiking threshold, *V_j_* ≥ *V_s_*, the voltage is reset with *V_j_* ← *V_r_*, clamped for the refractory time *T_ref_*, and the spike-triggered adaptation increment, *b*, is added to the adaptation current, *I_A,j_* ← *I_A,j_* + *b*.

The synaptic currents are described by a sum of excitatory and inhibitory contributions as

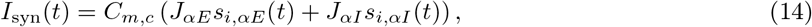

with the coupling strength *J_αβ_* from population *β* to *α*, and the fraction of active synapses *s_i,αβ_* ∈ [0, 1]. The synaptic dynamics is then given by

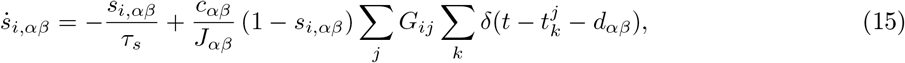

where *G_ij_* is a random binary connectivity matrix. The first sum is over all afferent neurons *j*, while the second sum is over all incoming spikes *k* from neuron *j* emitted at time *t_k_* after a delay *d_αβ_*. The (1 − *s_i,αβ_*) term in Eq. (15) acts as a saturation term and integrates all incoming spikes only if *s_i,αβ_* < 1, i.e., only if there is a synaptic capacity available.

In the mean-field approximation, the distribution *p*(*V*) of the membrane potentials, and the mean population firing rate *r* can be calculated using the Fokker-Planck equation in the thermodynamic limit, with the number of neurons *N* → ∞ [53, 43]. However, using the model reduction scheme described in [54, 43], we exploit the low-dimensional linear-nonlinear cascade model, where for a given mean membrane current *μ_α_* with standard deviation *σ_α_*, the mean 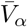 of the membrane potentials, the adaptive timescale *τ_α_*, and the population firing rate *r_α_* in the steady-state can be captured by a set of nonlinear transfer functions Φ(*μ_α_, *σ*_α_*) [55], i.e.

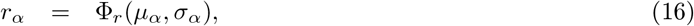

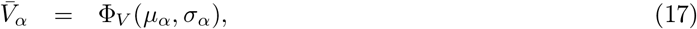

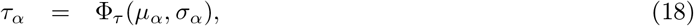

for *α* ∈ {*E, I*}. In the case of excitatory population (*α* = *E*), the mean adaptation current 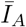 is subtracted from the mean membrane current *μ_E_* in the computation of population firing rate, mean membrane voltage, and adaptive timescale via transfer functions, i.e., 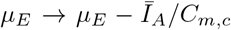. These transfer functions can be precomputed for a specific set of single AdEx neuron parameters [54] and are shown in Fig. A.1.

The average population currents in the mean-field approximation are given by

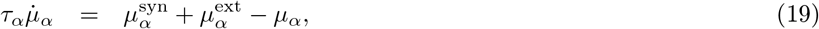

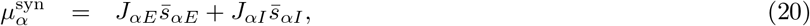

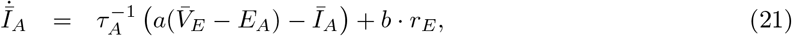

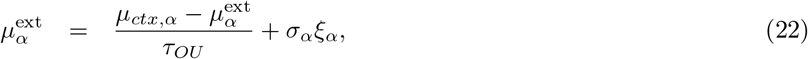

where the dynamics of mean membrane current *μ_α_* depends on the synaptic current 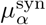 and an external noisy current 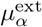, which enters the system in the form of Ornstein–Uhlenbeck process with mean drift *μ_α_*, standard deviation *σ_α_*, and time scale *τ_OU_*. *ξ_α_* is drawn from random Gaussian white noise process with zero mean and unit variance. In the subsequent text, we refer to mean drifts as *μ_E_* (*μ_I_*) as an input to the excitatory (inhibitory) population with the original physical units of mV/ms. After multiplying *μ_α_* with membrane conductance *C*, we obtain input currents in physical units of A, but we will omit the multiplication by *C* in subsequent text and treat all *μ_α_* with units of A.

The mean of the fraction of active synapses 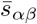 obeys

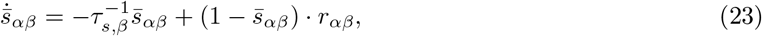

where the mean *r_αβ_* and the variance *ρ_αβ_* of the effective input rate from population *β* to population *α* for a spike transmission delay *d_αβ_* are given by

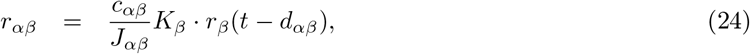

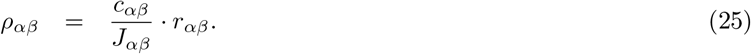

Finally, the current variance 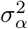, and the variance of the fraction of active synapses 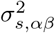 are given by

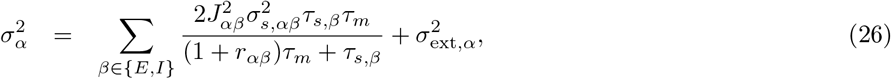

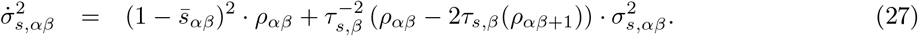

The parameters are summarized in Table in G.2.

### 2.4. Connecting the thalamic and cortical models

The thalamic and cortical models introduced before are coupled into a thalamocortical model with feedback and feedforward connections via their firing rate. More concretely, excitatory firing rate from cortical node enters the dynamics of both TCR and TRN populations in the thalamus with a connection strength *N_ctx→thal_* (cf. Fig. 1). In particular, the excitatory cortical firing rate comes in Eq. 8 for the excitatory AMPA synapse with *N_kk′_* = *N_ctx→thal_* and *r_k′_* = *r_E_*(*t* − *d_ctx,thal_*), with *d_ctx,thal_* being the thalamocortical delay of 13 ms.

For the thalamocortical connection, we connect excitatory firing rate from TCR population onto excitatory population of the cortical model (cf. Fig. 1). In particular, the TCR firing rate enters the cortical dynamics in Eq. 24 with *r_β_* = *N_thal→ctx_* · *r_TCR_*, and *d_αβ_* = *d_ctx,thal_*. The transmission delay in thalamocortical direction is set to the same value as in the corticothalamic direction, i.e., 13 ms.

### 2.5. Numerical simulations

The whole delayed dynamical equations system (Eqs. (1)–(27)) was integrated with the forward Euler method. If not mentioned otherwise, simulated time was *t* = 30 seconds with an integration timestep of *dt* = 0.01 ms. After integration, time series were subsampled at *dt_samp_* = 10 ms. The thalamocortical model was simulated using the `neurolib` library [56]. `neurolib` is a computational framework for whole-brain modeling written in `Python`. It provides a set of neural mass models and is designed to be extendable, and allows for easy implementation of custom mass models. Moreover, it supports heterogeneous brain modeling by coupling more than one type of mass model together. It offers a custom parameter exploration and optimization module for fitting models to multimodal experimental data using evolutionary algorithms. All subsequent analyses were also done in `Python`, and the repository with the model and analysis code is available at https://github.com/jajcayn/thalamocortical_model_study.

Background noise inputs are implemented as an independent Ornstein–Uhlenbeck processes [57] satisfying

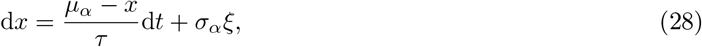

with the mean drift *μ_α_* and standard deviation *σ_α_*, for *α* ∈ {*E, I, TCR*}, integration timestep *dt*. *ξ* is drawn from random Gaussian white noise process with zero mean and unit variance. The Ornstein-Uhlenbeck processes, *x*, were pre-integrated and then inserted into the thalamic state equations as *ϕ′* into eq. (8), and to the cortical equations as 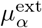 into Eq. (22).

### 2.6. Spindle detection from the model output

For automated spindle detection from model output, we used a modified version of the A7 spindle detection algorithm described in Lacourse et al. [58] and implemented in the `YASA` (*Yet Another Spindle Algorithm*) package [59] for `Python`.

Since the A7 spindle detection algorithm is designed to analyze empirical data, e.g., EEG, MEG, ECoG, or LFP (cf. [60, 61]), we made adjustments to the algorithm parameters. For detecting spindles on the cortical model output, we lowered the threshold for the duration from 0.5 to 0.3 seconds and for the relative power in the *σ*-band from 0.2 to 0.15.

### 2.7. Cross-frequency coupling (CFC) measures

In order to quantify the coupling between the phase of the slow oscillation and the spindle amplitude, we compute the Kullback-Leibler modulation index (*KL-MI*, cf. [62]) and the mean vector length (*MVL*, cf. [63]). For quantifying the phase-phase CFC, we use the phase-locking value (*PLV*, cf. [64]) and the mutual information (*MI*, cf. [65, 66]) between time series of instantaneous phases. We first filter the modeled outputs using a one-pass, zero-phase, non-causal finite impulse response (FIR) bandpass filter, implemented in the `mne Python` package [67] (either adapted to the SO, 0.1–3.0 Hz, or the fast spindle frequency, 12–15 Hz, ranges). The filtered signal is then passed to the Hilbert transform, which provides us with the estimate of the complex analytic signal 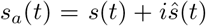, where *s*(*t*) is the real-valued input signal (model output), and 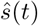 is the Hilbert-transformed signal. The instantaneous phase, *ϕ*(*t*), and amplitude, *A*(*t*), are then given by

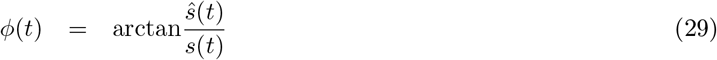

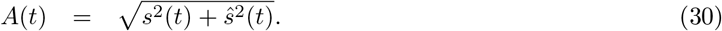

The *KL-MI* estimates the phase-amplitude coupling by computing the Kullback-Leibler divergence [68] between the distribution of spindle amplitudes *A_spindle_*(*t*) over the slow oscillation phase bins *ϕ_SO_*(*t*) and a uniform distribution (the null hypothesis of no phase-amplitude coupling). The *KL-MI* is defined by

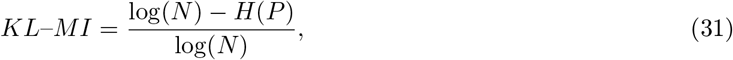

where *N* is the number of phase bins, and *H*(*P*) is the Shannon entropy of the amplitude distribution.

The MVL is computed by averaging the complex time series constructed by multiplying the spindle amplitude, *A_spindle_*(*t*), with the term containing the phase, *ϕ_SO_*, of the slow oscillations, thus

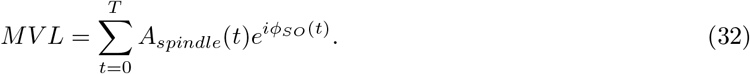

The length (real part) of the MVL vector quantifies the strength of phase-amplitude coupling. Its phase denotes the mean phase of the slow oscillation at which the spindle amplitude is strongest.

The PLV, as a measure of phase locking between two oscillations, is computed by temporally averaging the phase differences on unit circle between the phase of the slow oscillation and the phase of the spindle oscillation:

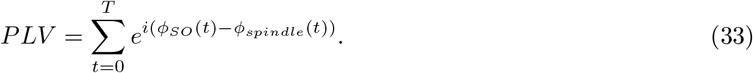

As with the MVL, the length of PLV vector indicates the strength of phase-locking, while the phase represents the phase shift.

Finally, the *MI* of two discrete variables (in our case, the time series of estimated phases) can be computed using

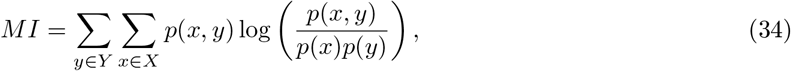

where *p*(*x, y*) signifies the joint probability mass function of {*X, Y*}, and *p*(*x*) and *p*(*y*) represents the marginal probability mass functions of *X* and *Y*, respectively. All three probability mass functions are estimated using an equiquantal binning algorithm [69] with 16 bins.

To test for statistical significance, we computed 1000 Iterative Amplitude Adjusted Fourier Transform (IAAFT) surrogates from a randomization procedure that preserves the power spectrum and the amplitude distribution of the original time series [70]. The surrogates were constructed by iterative replacements of Fourier amplitudes with the values from the original time series and by rescaling the distribution match between the distribution and the power spectrum of the original data. We opted to use IAAFT surrogates since our modeled data are firing rates with a non-Gaussian distribution, and do not meet the criteria for using the basic Fourier Transform surrogates. After constructing the surrogates, we computed a given CFC measure for each surrogate time series, yielding an empirical null distribution. Finally, we compared a result on the modeled output to this null distribution to obtain the empirical p-value.

## 3. Results

### 3.1. Dynamical repertoire of the thalamic model — spindle oscillations

The isolated thalamic node model generates spindle oscillations (see Fig. 2). As described previously [34, 71, 72], spindle oscillations emerge through the reciprocal interaction of the TRN, which acts as a pacemaker, and the TCR, which mediates spindle propagation to the cortex [19, 11]. A low-threshold burst discharge in the TRN population causes synchronous and robust inhibition of the TCR, which activates its T-type calcium current. The subsequent activity rebound drives the TRN population to elicit additional low-threshold bursts. Additionally, activating the T-type calcium current requires a strong tonic hyperpolarization caused by the potassium leak current [51, 71].

The waxing and waning structure of the spindle oscillations is caused by the afterdepolarization in the TCR, mediated by the hyperpolarization-activated cation-nonselective channels, represented by the *I_h_* current [11]. A sequence of low-threshold spikes leads to the build-up of calcium in the TCR cells, which increases the effective conductivity *g_h_* of *I_h_*. The depolarization of the TRN additionally counteracts its ability to produce a low-threshold spike, which conclusively ceases the spindle oscillation [73, 30]. Sleep spindle termination also involves cortical and brain stem mechanisms [11], which we omit for brevity here. The two mechanisms — one creating fast spindle oscillation between 12–15 Hz, the other responsible for the waxing and waning structure with frequency < 1 Hz — are visualized in Fig. 2.

We probed the thalamic model for a set of parameters that convey the generation of spontaneous spindles. Fig. 3 summarizes spindle activity in the thalamic model as a function of three parameters: the conductances of the potassium leak, *g_LK_*, and rectifying, *g_h_*, currents, and the variance, *σ_TCR_*, of the background noise input. In the noise-free case (Fig. 3A, *σ_TCR_* = 0.0 mV/ms^3/2^), we observe two regions where spindle oscillations emerge spontaneously (marked as region I and region II in the figure). In these two regions, the interaction between the fast T-type current and its slow modulation via the rectifying current leads to spindle oscillations. The regions differ in the value of *g_LK_* (cf. [34]). Spindle oscillations in the region I are longer, more symmetric, and have a longer inter-spindle interval (Fig. 3B rows I vs. II).

**Figure 3:**
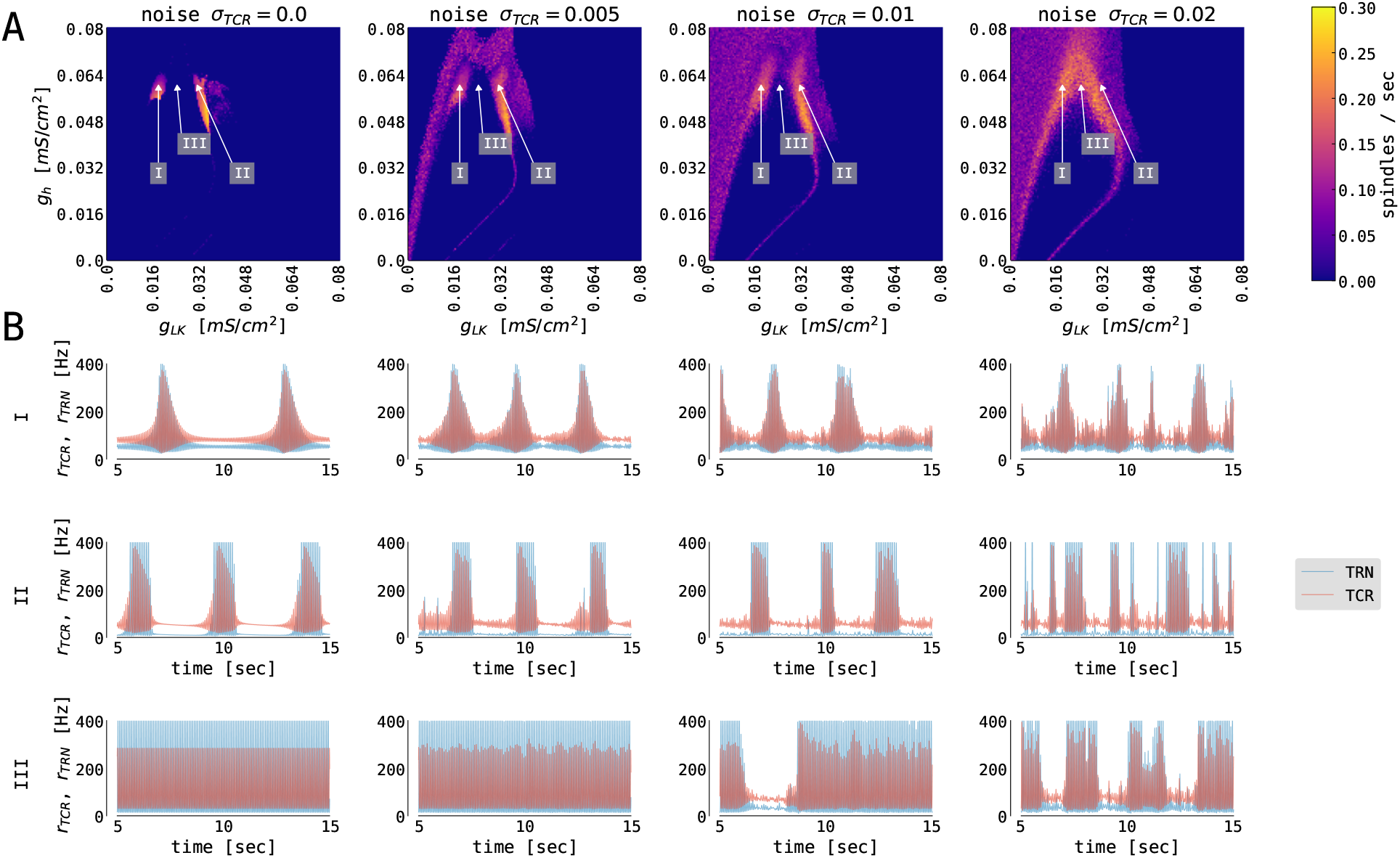
The generation of thalamic sleep spindles depends on the conductances *g_LK_* and *g_h_*. (**A**) Color-coded number of spindles per second as a function of conductance parameters *g_h_* and *g_LK_* for 4 different noise levels *σ_TCR_*. The model was simulated for 65 seconds. The first 5 seconds were not included in the statistics. (**B**) 10 seconds time series from the thalamic node model (red traces for TCR firing rate, blue traces for TRN firing rate) from the three regions marked in (**A**) with different parameters: *g_LK_* = 0.018 mS/cm^2^ (I), *g_LK_* = 0.031 mS/cm^2^ (II), *g_LK_* = 0.024 mS/cm^2^ (III), and *g_h_* = 0.062 mS/cm^2^ (all three regions). For other model parameters, see Table G.1.

When we introduce noise to the thalamic model (second to fourth column of Fig. 3), we observe qualitative changes in the spindling behavior. The most noticeable is the emergence of the waxing and waning cycle as a function of noise strength in areas neighboring the original spindle-promoting regions. This is clearly seen in the third row of Fig. 3B (region III), as the spindles emerge for sufficiently strong background noise. In the spindle-promoting regions I and II, noise randomizes spindle timing: it can push the TCR population into a spindle event or, reversely, cease spindling faster than the slow *I_h_*-driven negative feedback loop alone. The background noise in our model can act as a depolarizing or hyperpolarizing force, thus can either speed up the activation of *I_h_* and cease spindling, or, reversely, slow it down and prolong the spindling periods. Sufficiently strong noise may also change the number of spindles in the spindle-promoting regions I and II as seen for noise strengths *σ_TCR_* = 0.01 mV/ms^3/2^ and larger.

To the best of our knowledge, there is no experimental evidence of how spindle properties change as a function of *g_LK_*. However, we know that spindle density ranges between 2–10 spindles/min in EEG data [75, 11], and spindles can have various levels of symmetry. Although, there seems to be a continuous range of spindle density and symmetry rather than two clusters in the conductance phase space. We consider the two separated spindling regions as a pure model feature (which is confirmed by the bifurcation analysis of the thalamic model in [34], Fig. 2), where the model can generate more symmetrical and less frequent spindles (region I, lower *g_LK_*, hence lower hyperpolarization), and less symmetrical and more frequent spindles (region II, higher *g_LK_*).

Outside of the spindle-promoting regions I and II, the thalamic model displays continuous oscillation in the *σ*-band due to hyperpolarization induced rebound bursts (as in region III Fig. 3B, but also for larger and smaller values of *g_h_*, hence above and below regions I and II). This behavior does not exhibit the waxing and waning structure since the T-type current dominates and *I_h_* is not strong enough to sufficiently depolarize the TRN population to cease the oscillation. Furthermore, for larger values of *g_LK_* (to the right of region II in Fig. 3) the *I_LK_* current dominates, and the thalamic model switches to slow, *δ*-like rate oscillations. In the corners of the state space spanned by the conductances *g_LK_* and *g_h_* the thalamic model exhibits a stable fixed point behavior with constant firing rates (e.g. for maximal conductances *g_h_* = *g_LK_* = 0.08 mS/cm^2^ the TCR exhibit down state with constant rate of 10 Hz, while for *g_h_* = *g_LK_* = 0 mS/cm^2^ the TCR exhibit up state with constant rate of 116 Hz).

In its default parametrization, our model exhibits a spindle frequency of approximately 13 Hz. The spindle frequency in the thalamic model depends on the conductance of the T-type calcium current, *g_T_* [34]. By changing the conductance value in the TCR population, the model is able to reproduce spindle frequencies in the whole range of fast spindle oscillations (approximately 11.5 – 15 Hz). On the other hand, changing the value of *g_T_* in the TRN population has only minor effect on the spindle frequency. Significantly, this effect does not qualitatively change with the introduction of noise into the thalamic model. Other spindle parameters, such as its duration, amplitude, or symmetry, are approximately invariant with respect to changes in *g_T_* in both populations.

### 3.2. Dynamical repertoire of the cortical model — slow oscillations

The cortical node as a motif of delay-coupled excitatory and inhibitory populations with somatic spike-frequency adaptation can be parametrized into four different dynamical regimes dependent on the mean external input currents *μ_α_* to both populations *α* ∈ {*E, I*} [43]. The 2D slices of bifurcation diagrams of interest are shown in Fig. 4A. When the excitatory input is weak, the system is in its DOWN state: a fixed point solution with a low firing rate of the excitatory population. As we increase the external input to the E population, the system undergoes a supercritical Hopf bifurcation, and a limit cycle emerges. For a weak background current to the I population, the interaction between inhibition and excitation creates an E-I oscillation with a frequency of ~25 Hz. For stronger background current to the I population, the interplay between adaptation and excitation gives rise to a slow limit cycle with frequencies of around 2 Hz and lower. The system is in a stable fixed point for stronger background current to the E population again, albeit with a higher firing rate, the so-called UP state [43].

**Figure 4:**
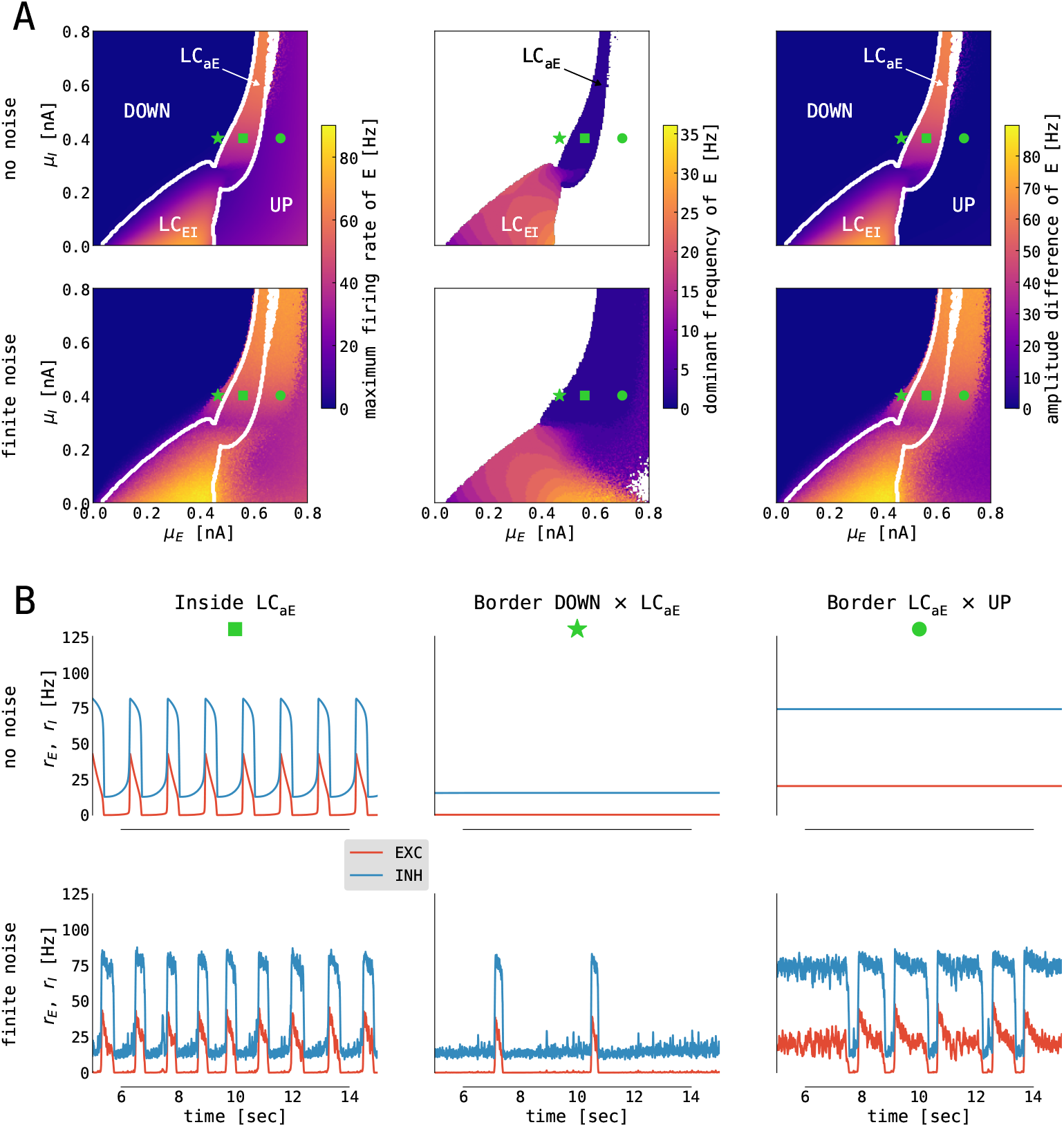
Emergence of slow oscillations in the cortical model. (**A**) Slices of state space spanned by the mean external input currents *μ_α_*, *α* ∈ {*E, I*}, to the excitatory and inhibitory population with no (*σ_E_* = *σ_I_* = 0.0 mV/ms^3/2^, top panel) and finite (*σ_E_* = *σ_I_* = 0.05 mV/ms^3/2^, bottom panel) noise. The panels show (left to right): maximum firing rate of the excitatory population, dominant frequency of its rate oscillations (computed as a frequency with maximum power in its Welch spectrum; white color denotes no oscillations, i.e. fixed point dynamics), and the difference between its maximum and minimum firing rates. White lines indicate boundaries between fixed points (*UP* and *DOWN*) and limit cycles (*LC_EI_* and *LC_aE_*) computed on the noise-free case. (**B**) Simulated time series (from E in red, from I in blue) from selected states for no (*σ_E_* = *σ_I_* = 0.0 mV/ms^3/2^, top panel) and finite (*σ_E_* = *σ_I_* = 0.05 mV/ms^3/2^, bottom panel) noise. Panels show (left to right) traces obtained from inside the limit cycle (*μ_E_* = 0.56 nA, *μ_I_* = 0.4 nA, marked with green square in **A** panels), from the left (*μ_E_* = 0.466 nA, *μ_I_* = 0.4 nA, marked with green star in **A** panels), and the right (*μ_E_* = 0.7 nA, *μ_I_* = 0.4 nA, marked with green circle in **A** panels) border of the limit cycle. For other parameters, see Table G.2

When noise is added to a system parametrized close to the border of the slow limit cycle and the DOWN state, we observe a DOWN state with occasional irregular UP state excursions (i.e., noise pushes the system intermittently into the limit cycle — Fig. 4B middle). Reversely, along the border between the limit cycle and the UP state, the system exhibits a UP state with irregular DOWN state excursions caused by noise (Fig. 4B right). From the perspective of modeling slow oscillation during deep sleep, the optimal operating point of the cortical model is close to the border between the UP state and the slow limit cycle, where the cortical node undergoes irregular DOWN state excursions [76].

### 3.3. Thalamic model driven by cortical input

We first investigate the state space of the thalamic model by studying the effects of external (cortical) inputs without closing the feedback loop back to the cortex. Fig. 5 shows the number of thalamic spindles as a function of the potassium leak conductance *g_LK_* and the conductance *g_h_* of TCR’s rectifying current for constant external rate inputs to both thalamic populations. For larger external inputs, the regions of thalamic spindling translate to lower values of *g_h_* independent of TCR noise levels. Increased external input leads to a higher concentration of intracellular calcium in the TCR, which activates the rectifying *I_h_* current more strongly and ultimately abolishes the waxing and waning cycle of spindle oscillations (cf. Fig. 2). For smaller values of *g_h_, I_h_* is reduced, and the waxing and waning of spindle oscillations reappears.

**Figure 5:**
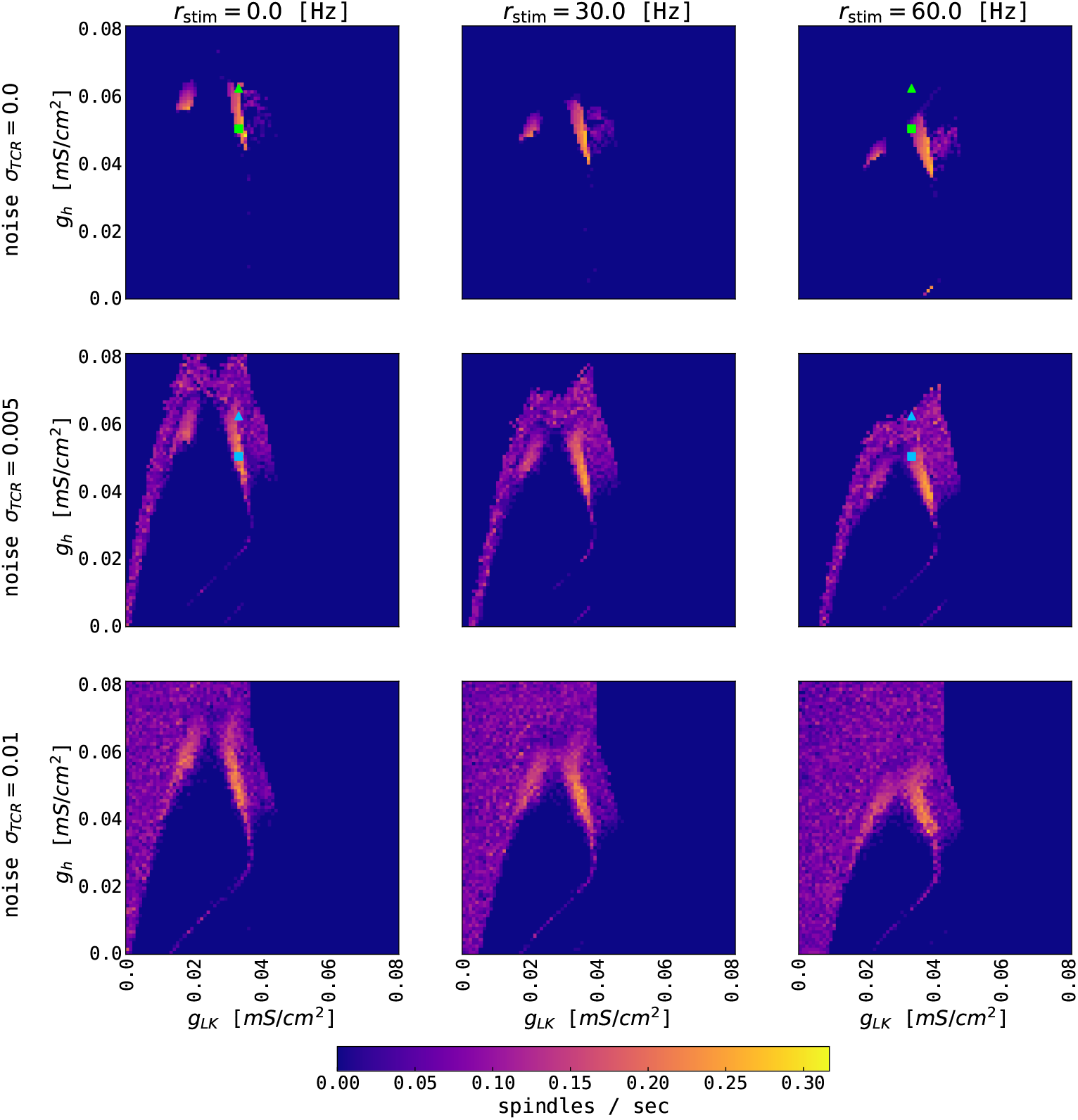
Effects of constant external input on thalamic spindling. The panels show the color-coded rate of thalamic spindles as a function of the conductance parameters *g_h_* and *g_LK_* for three different noise levels *σ_TCR_* and three different strengths *r_stim_* of the external input. External coupling strengths are equal for the TCR and TRN populations and are set to one. The other model parameters of the thalamic node are given in Table G.1. Squares (*g_h_* = 0.05 mS/cm^2^, *g_LK_* = 0.033 mS/cm^2^) and triangles (*g_h_* = 0.062 mS/cm^2^, *g_LK_* = 0.033 mS/cm^2^) mark parameter values used for Fig. 6. Symbol color refers to the different thalamic noise levels. Note that the typical excitatory firing rate of the cortical node’s UP state is up to 50 – 60 Hz.

We then stimulate the thalamic node with rectangular current pulses mimicking an idealized sequence of cortical UP and DOWN states. Fig. 6A shows the resulting rates of the TCR and TRN populations as a function of time. For higher values of *g_h_*, thalamic spindles are only induced in the DOWN state, as already expected from Fig. 5. For lower values of *g_h_*, spindling resumes during UP states, albeit with lower spindle event frequency. Finally, we stimulate the thalamic node with the excitatory rate of a cortical node undergoing sustained adaptation-induced slow oscillations (Fig. 6B) and noise-induced DOWN-state transitions (Fig. 6C). With few exceptions, thalamic spindles are induced right after cortical UP to DOWN transitions, but — given the short duration of the cortical DOWN state — every SO induces one thalamic spindle only. In this setting, this motif well reproduces the SO-correlated spindling observed during NREM sleep [9, 40, 41, 42]. Changing the conductance *g_h_* of TCR’s rectifying current, the ratio between the numbers of “free” and SO-correlated spindle events can be adapted.

**Figure 6:**
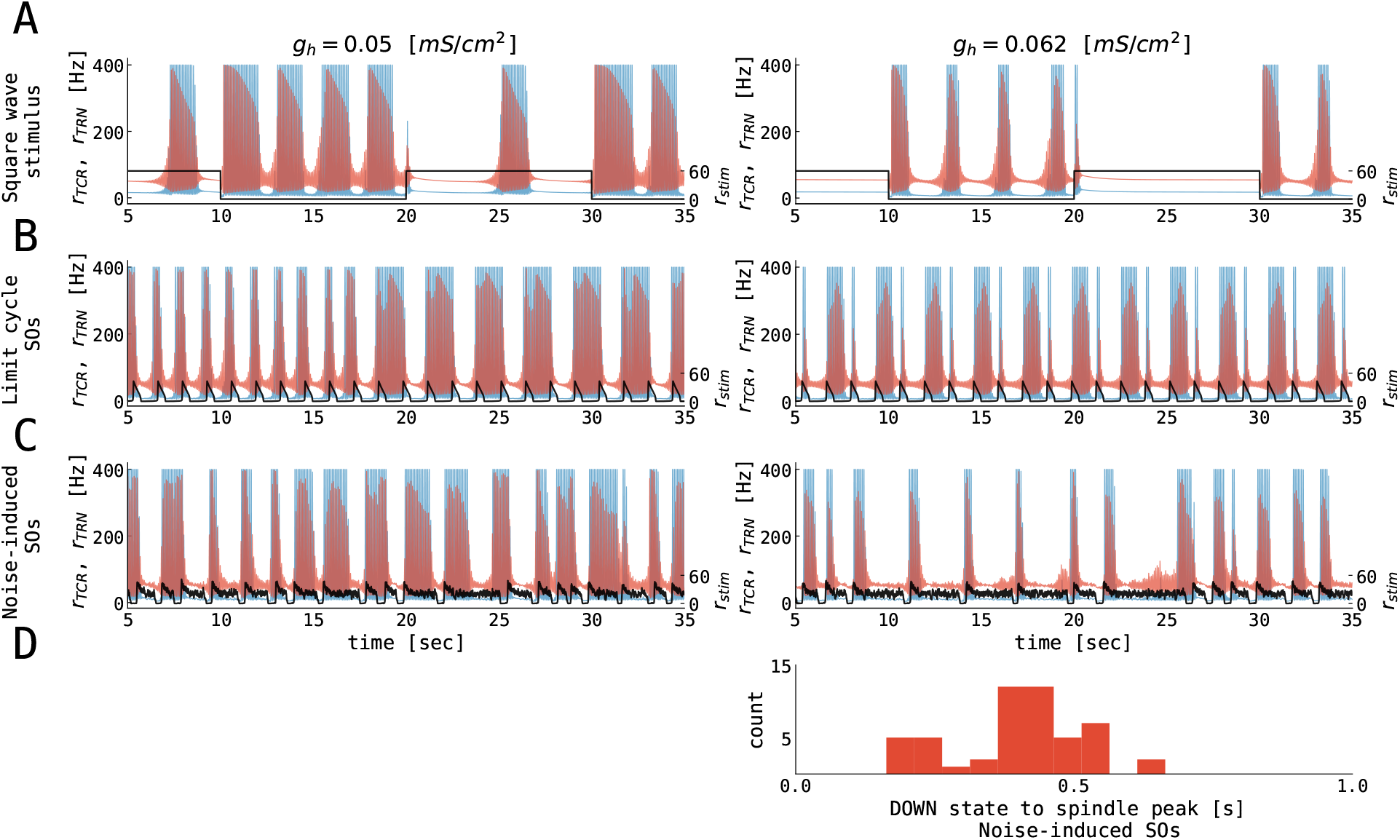
Thalamic response to alternating cortical UP and DOWN states. The (**A**) – (**C**) panels show the firing rates of the TCR (red), the TRN (blue), and the cortical input (black) as a function of time. Corticothalamic coupling strength (*N_ctx→thal_* = 1.0) was equal for the TCR and TRN populations. (**A**) Thalamic response to square wave stimulation of 0.05 Hz with the amplitude of 60 Hz. (**B**) Thalamic response to excitatory rate input, which is generated by a cortical node undergoing limit cycle oscillations (cf. Fig. 4B, upper left panel). (**C**) Thalamic response to excitatory rate input, which is generated by a cortical node undergoing noise-induced DOWN state transitions (cf. Fig. 4B, lower right panel). (**D**) Distribution of delays between midpoints of cortical DOWN states and spindle-band peaks in the noise-induced slow oscillations from panel (**C**). The parameters for the thalamic node were: *g_h_* = 0.05 mS/cm^2^ (left panels, see also green squares in Fig. 5), *g_h_* = 0.062 mS/cm^2^ (right panels, see also green triangles in Fig. 5), *g_LK_* = 0.033 mS/cm^2^, for all other parameters see Table G.1. The parameters for the cortical node were: *μ_E_* = 0.56 nA, *σ_E_* = *σ_I_* = 0.0 mV/ms^3/2^ for panels (**B**) and *μ_E_* = 0.7 nA, *σ_E_* = *σ_I_* = 0.05 mV/ms^3/2^ for panels (**C**), for all other parameters see Table G.2. The parameters of connected thalamocortical model are given in Table G.3.

### 3.4. Cortical model driven by thalamic input

Secondly, we study the effects of external thalamic inputs to the cortical node with a one-way thalamus → cortex connection. Thalamic input (noise-free spindles) was simulated using our thalamic model parametrized in the spindling region I (Fig. 3), i.e., with *g_LK_* = 0.018 mS/cm^2^, *g_h_* = 0.062 mS/cm^2^. Fig. 7 summarizes cortical responses as a function of cortical parameters and thalamus → cortex connection strengths. We observe two distinct effects of thalamic stimulation. Firstly, it increases the slow oscillation frequency because additional thalamic input leads to an increase of the excitation, an effect similar to increasing *μ_E_* in the isolated cortical node.

**Figure 7:**
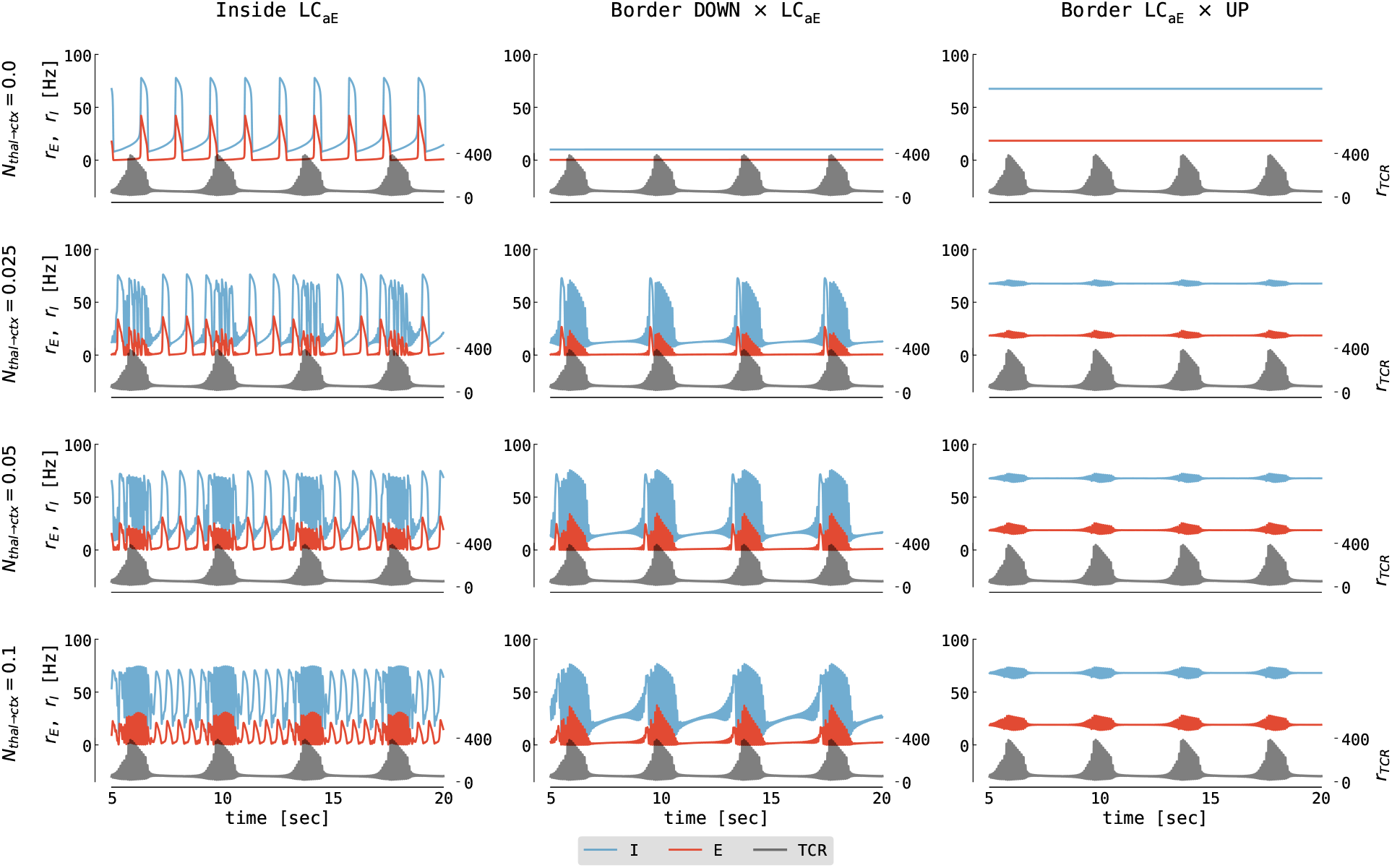
Cortical response to thalamic spindle stimulation. The panels show the firing rates of the excitatory cortical population (red), the inhibitory cortical population (blue), and the thalamic input (black) as a function of time. Columns show different parametrizations of the cortical node (left to right): inside the slow limit cycle (*μ_E_* = 0.56 nA, cf. Fig. 4B left), the border of the DOWN state and the limit cycle (*μ_E_* = 0.466 nA, cf. Fig. 4B middle), and the border of the limit cycle and the UP state (*μ_E_* = 0.7 nA, cf. Fig. 4B right). Different rows show different values of thalamus → cortex connection strength ranging from *N_thal→ctx_* = 0.0 to *N_thal→ctx_* = 0.1. The parameters for the cortical node were: *μ_I_* = 0.4 nA, *σ_E_* = *σ_I_* = 0.0 mV/ms^3/2^, for all other parameters see Table G.2. The parameters for thalamic node were: *g_LK_* = 0.018 mS/cm^2^, *g_h_* = 0.062 mS/cm^2^, *σ_TCR_* = 0.0 mV/ms^3/2^, for all other parameters see Table G.1. The parameters of connected thalamocortical model are given in Table G.3.

The second effect is the imprinting of spindle oscillations into the cortical activity for stronger coupling strengths (Fig. 7, lower two rows). Typically, a spontaneous thalamic spindle induces a transition to a cortical UP state, and spindle activity is superimposed on the UP state of the cortical activity. After the thalamic spindle ceases, the thalamic firing rate significantly decreases, and the cortex returns to the DOWN state due to the adaptation mechanism of excitatory neurons.

The addition of noise to the cortical node (*σ_E_* = *σ_I_* = 0.05 mV/ms^3/2^) does not qualitatively change our previous observations. However, background noise can induce a DOWN swing (leading to irregular slow oscillation) in the cortex parametrized at the border between the limit cycle and the UP state (Fig. B.2). Moreover, increasing the thalamus → cortex connection strength leads to prolonged UP states, to less frequent DOWN states, and, for large enough coupling strength, to spindle activity from the thalamus being imprinted onto the UP state of the cortex.

### 3.5. Full thalamocortical loop

In the following section, we study the dynamics of the full thalamocortical loop consisting of one cortical and one thalamic node, coupled according to Fig. 1. To summarize our hypotheses regarding the thalamocortical motif: we expect slow oscillations to emerge in the cortex and spindles in the thalamus [6, 77]. Additionally, we expect slow oscillation activity to affect the timing of thalamic spindles [78, 79] and thalamic spindles to cause cortical spindles during cortical UP states [80, 41, 42].

Fig. 8 shows the amplitude difference and the power in two spectral bands for the firing rate of the cortical excitatory population, depending on the connection strengths of thalamus → cortex and cortex → thalamus. Increasing thalamus → cortex connection strength leads to a broader region in which the cortical node is oscillating in the slow oscillation band, albeit the amplitude of cortical oscillation decreases (Fig. 8A). Moreover, increasing the thalamus → cortex connection strength also causes a power decrease in the slow oscillation band (0.1–3.0 Hz), and the region of increased slow oscillation power shifts slightly towards lower background excitation values (to the left in the panels), as depicted in Fig. 8B. Finally, the power in the *σ*-band (11.0–16.0 Hz) increases as a function of both thalamus → cortex and cortex → thalamus connection strengths (Fig. 8C). Adding background noise to both cortical (*σ_E_* = *σ_I_* = 0.05 mV/ms^3/2^) and thalamic (*σ_TCR_* = 0.005 mV/ms^3/2^) nodes does not qualitatively change these observations (not shown). Thalamic parameterization was chosen to from the spindling region II (Fig. 3), i.e., the thalamus was simulated with *g_LK_* = 0.033 mS/cm^2^, *g_h_* = 0.062 mS/cm^2^.

**Figure 8:**
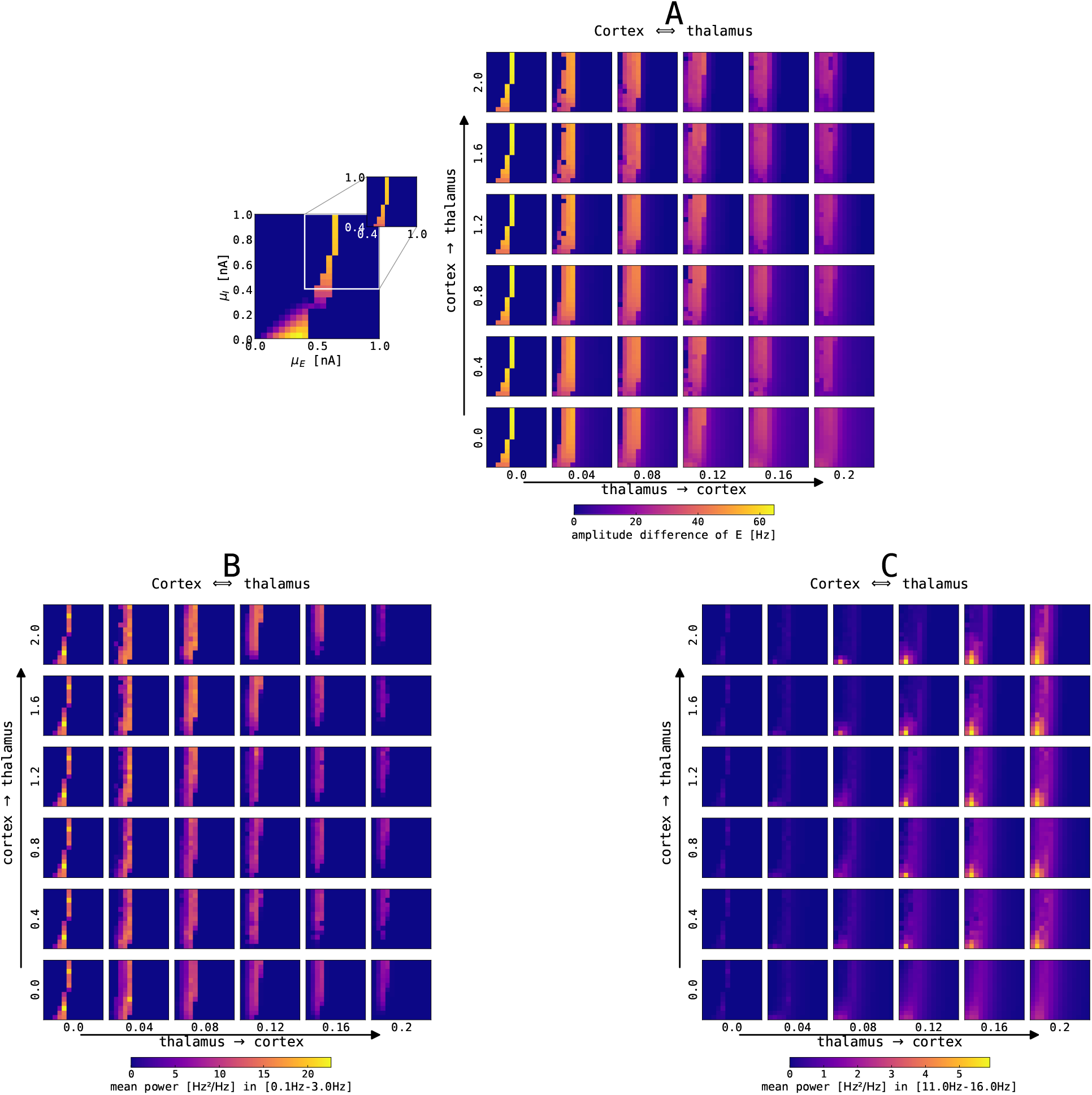
Impact of changes of thalamocortical connection strengths on cortical SO and spindle activity. The panels show a single slice of the state space diagram of the cortical model on par with Fig. 4, zoomed into the slow limit cycle region, for the recurrently connected thalamocortical model of Fig. 1. The full model was simulated with different strengths of connections in both directions: thalamus → cortex is shown on the x-axis, and cortex → thalamus is shown on the y-axis. Panels show (**A**) the color-coded difference between the maximum and minimum firing rate in the excitatory cortical population, (**B**) the color-coded mean spectral power in the slow oscillation range (0.1–3 Hz), and (**C**) the color-coded mean spectral power in the fast spindle range (11–16 Hz). The mean spectral power in both bands was computed by averaging the power spectral density computed using Welch’s method within the slow oscillation band, 0.1–3.0 Hz, and *σ* band, 11–16 Hz. Note that the bottom left panels (*N_ctx→thal_* = 0.0, *N_thal→ctx_* = 0.0) parallel the bifurcation diagram of isolated cortex in Fig. 4. All simulations were conducted without noise, for other parameters see Tables G.1, G.2 and G.3.

### 3.6. Cortical spindle activity in the thalamocortical loop model

Fig. 9A summarizes cortical spindle activity dependent on thalamus → cortex and cortex → thalamus connection strengths and the level of background excitation and inhibition (*μ_E_* and *μ_I_*, respectively). The density of cortical spindles is directly proportional to the thalamus → cortex connection strength. Cortical spindle activity emerges mainly in the region where cortical node exhibits oscillatory activity (cf. Fig. 8A).

**Figure 9:**
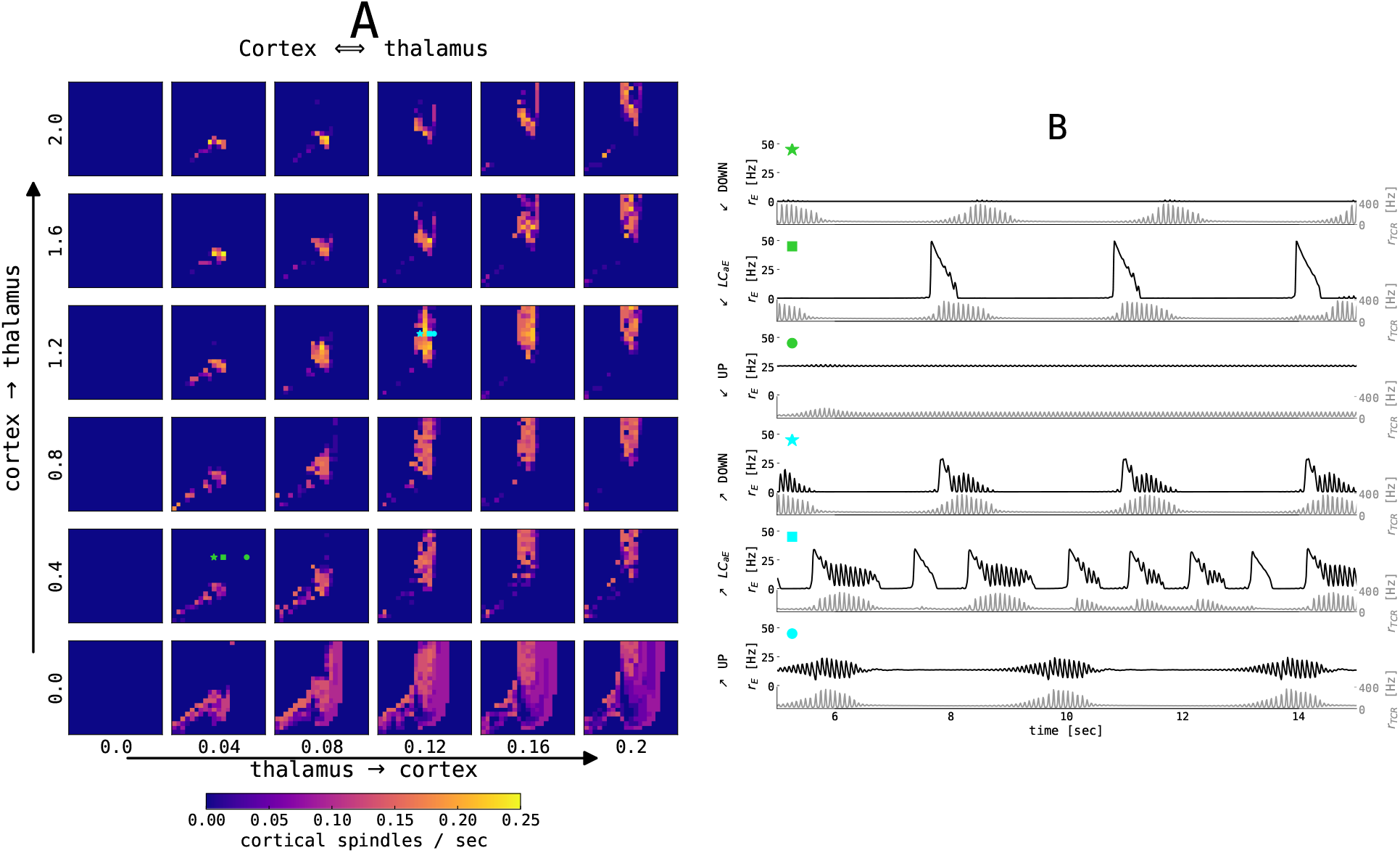
Interaction between cortical slow oscillation and thalamic spindles in the noise-free case. (**A**) Estimated number of cortical spindles per second as a function of the mean external input currents *μ^ext^* to the excitatory and inhibitory population (encoded by *x* and *y*-axis in each smaller panel, respectively) and the thalamus → cortex and cortex → thalamus connectivity strength (encoded by *x* and *y*-axis over panels, respectively). The thalamocortical model was simulated for 65 seconds without noise. (**B**) 15 seconds excerpts of time series from the excitatory population in the cortical node (black) and TCR in the thalamic node (gray) for two sets of connection strengths (*N_thal→ctx_* = 0.04 and *N_ctx→thal_* = 0.4, denoted by lime green symbols; and *N_thal→ctx_* = 0.12 and *N_ctx→thal_* = 1.2, denoted by aqua blue symbols) and three values of *μ_E_* as per three dynamical states of the cortex: DOWN state with *μ_E_* = 0.45 nA (star), slow limit cycle with *μ_E_* = 0.55 nA(square), and UP state with *μ_E_* = 0.8 nA for weaker connection strengths and *μ_E_* = 0.6 nA for stronger connection strength (circle). The points of interest are also denoted in (**A**). For all points *μ_I_* = 0.7 nA, and other parameters were kept constant as per tables G.1, G.2, and G.3.

The time series of firing rates in both nodes in the thalamocortical model is presented in Fig. 9B. For selected values of model parameters, at the border between the cortical DOWN state and the limit cycle, a spontaneous thalamic spindle causes the cortical node to go into the UP state, and the cortical activity is then shaped by the incoming spindle, depending on the thalamus → cortex connection strength as seen in Fig. 9B (aqua blue and lime green star). In the slow limit cycle, cortical parametrization for the higher connection strengths, the cortical UP states, and thalamic spindle waxing become coupled (Fig. 9B, aqua blue and lime green square). Typically, the thalamic node elicits a spindle after cortical UP to DOWN state transition, which creates a thalamic DOWN state. This allows the thalamic node to go into hyperpolarization and generate a spindle.

The cortical UP state activity is slightly modulated by the spindle oscillations of the thalamic node (Fig. 9B, lime green and aqua blue circles). With the addition of noise, this regime becomes interesting for our investigation, as seen in the next section.

### 3.7. Slow oscillation–spindle interaction in the UP state regime

In the UP state-dominant regime, the cortical model is parametrized in the UP state, with irregular DOWN swings caused by the background noise pushing the state into the limit cycle. The average length of the UP state depends on the amount *μ_E_* of background excitation.

For *μ_E_* = 0.61 nA, i.e., close to the bifurcation, UP states are of shorter duration (see Fig. 10). In this case, the UP and DOWN states are relatively regular, with cortical DOWN states providing a window of opportunity for hyperpolarization of the TCR and subsequent spindle generation, which then embed spindle oscillatory activity in the cortical node typically just after the DOWN to UP state transition. This behavior can be observed by following vertical white dashed lines in the cortical time series (Fig. 10A), which denote the midpoints of cortical DOWN states which are followed by a spindle within a 1.5 second window. These lines extend to the time-frequency representation, and we can see an intermittent increase of the cortical power in the spindle band, thus demonstrating cortical spindles nested in the UP states. The transition from the DOWN to the UP state in the cortical node is accompanied by an overshoot in the cortical activity, which then decreases due to the adaptation current, as seen in Fig. 10B. The mechanistic explanation of the SO–spindle relationship is illustrated by cortical DOWN-state and thalamic spindle time-locked plots (Fig. 10B and D): cortical DOWN state is followed by a waxing period in the thalamic activity, the spindle activity is then imprinted onto cortical activity, followed by an approximately 1 second long cortical spindle. Thus, spindles are typically observed during the peak of cortical UP state or shortly after, see circular histograms in Fig. 10B and D. Our model results align well with the data-driven studies [41, 42]. Mechanistically, a sustained UP state in the cortex suppresses spindling via the corticothalamic connections, since TCR cannot reach necessary hyperpolarization. Therefore, for this particular parametrization (predominant cortical UP states with irregular DOWN swings), we conclude the causal direction goes in the sense of cortex → thalamus.

**Figure 10:**
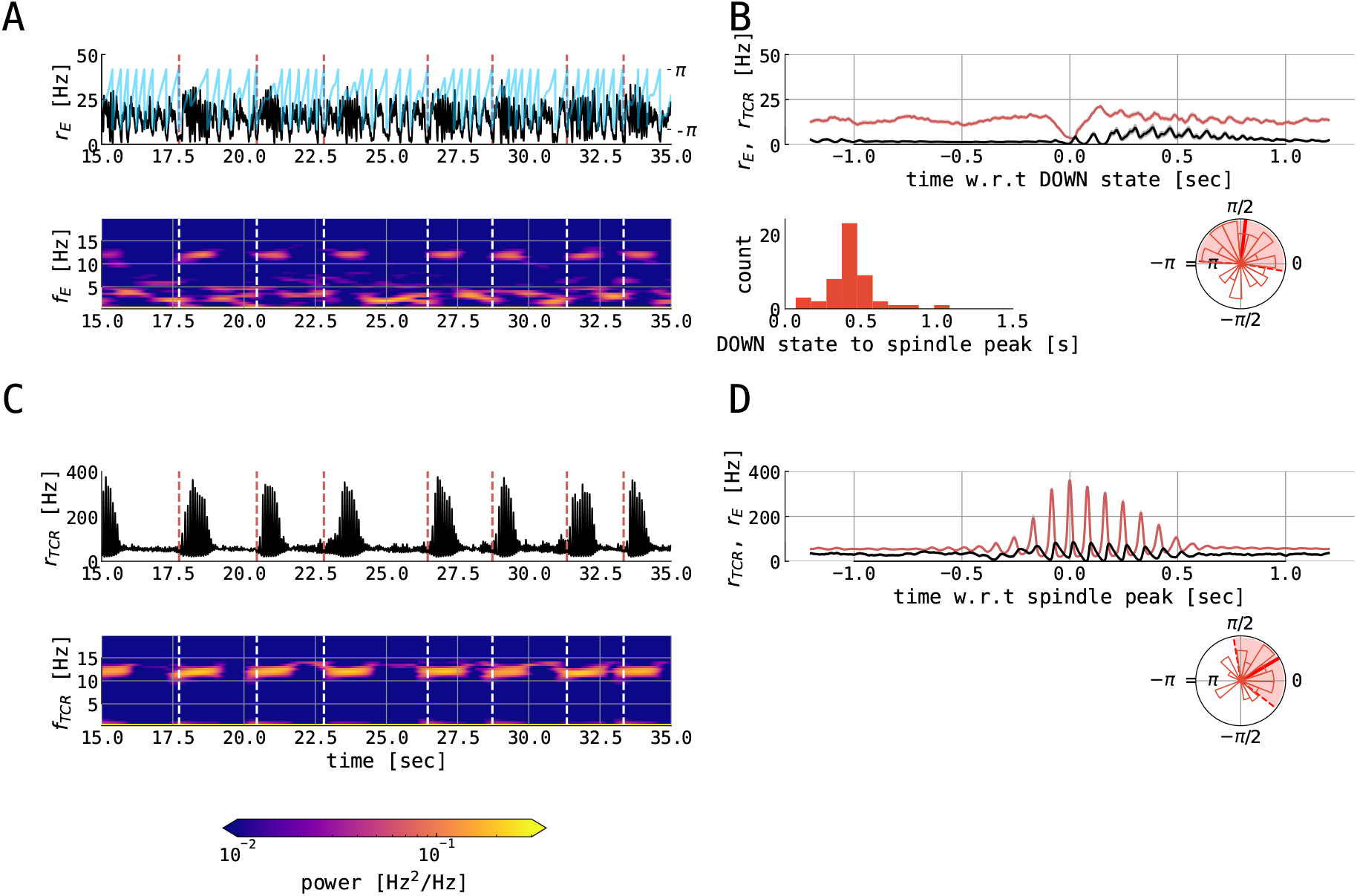
Spindles in the thalamocortical motif with short UP states. The figure shows various variables from 120 second of simulation of full thalamocortical model simulation with the cortical node being parametrized at the right border between the slow limit cycle and the UP state, relatively close to the bifurcation line (*μ_E_* = 0.61 nA). Individual panels show: (**A**) 20 seconds time series excerpt of the firing rate of the excitatory population in the cortical node (black) and its slow oscillation phase (blue) computed using the Hilbert transform of low-pass filtered cortical excitatory firing rate. The panel below shows the time-frequency representation of the cortical excitatory firing rates computed using the Short Time Fourier Transform with a 2 second time window. Dashed vertical lines (red in time series plot and white in time-frequency plot) denote the midpoints of cortical DOWN states which are followed by a cortical spindle in the 1.5 second window. (**B**) the mean ± the SEM of the cortical excitatory firing rates (red) and TCR firing rates (black), locked on the cortical DOWN states for the whole interval of 120 seconds. The panel below shows the distribution of cortical slow oscillation phases (shown in (**A**) with blue) for the maximum of the cortical *σ*-band peak. Shown are histogram bars in thin red and circular mean (± circular STD) in thick (dashed) red. (**C**) 20 seconds time series excerpt of the firing rate of the thalamocortical relay population in the thalamic node (black). The panel below shows the time-frequency representation of the TCR firing rates computed using the Short Time Fourier Transform with a 2 second time window. Dashed vertical lines (red in time series plot and white in time-frequency plot) denote the midpoints of cortical DOWN states which are followed by a cortical spindle in the 1.5 second window. (**D**) the mean ± the SEM of the TCR firing rates (red) and cortical excitatory firing rates (black), locked on the thalamic spindle peaks in the whole interval of 120 seconds. The panels below show: the distribution of delays between midpoints of cortical DOWN states and spindle-band peaks; and the distribution of cortical slow oscillation phases for the maximum of the thalamic *σ*-band peak. Shown are histogram bars in thin red and circular mean (± circular STD) in thick (dashed) red. The model was simulated with *N_thal→ctx_* = 0.12, *N_ctx→thal_* = 1.2, *μ_I_* = 0.4 nA, *σ_E_* = *σ_I_* = 0.05 mV/ms^3/2^, *σ_TCR_* = 0.005 mV/ms^3/2^, while other parameters were kept constant as per tables G.1, G.2, and G.3.

For a parametrization further away from the bifurcation (*μ_E_* = 0.66 nA), prolonged UP states are observed (Fig. C.3). The SO–spindle interactions are similar as in the case with shorter UP states (Fig. 10), however these prolonged UP states are more similar to what is seen during human NREM sleep [41, 42].

We also probed our connected thalamocortical model in the remaining two cortical parametrizations. In particular, when the cortex is parametrized in the DOWN state (with *μ_E_* = 0.36 nA, other parameters unchanged with respect to previous text), the causal pathway between cortex and thalamus reverses its direction. In this case, the cortex is predominantly in the DOWN state. Without a sustained cortical drive, the thalamus can generate free spindles, and these spindles, in turn, drive the cortex into the UP state as they are projected (Fig. D.4). Finally, when the cortex is parametrized in the limit cycle (*μ_E_* = 0.42 nA), its UP states hinder the generation of free spindles in the thalamus, and the thalamic spindles are generated only on prolonged DOWN states. When the thalamus generates a spindle within the window of the cortical DOWN state, the projection of the spindle then forces the cortex to the DOWN state—UP state transition (Fig. E.5).

Following previous data-driven results on the slow wave–spindle nesting [44, 9, 41, 42], we finally quantified the phase-phase and phase-amplitude coupling between these two rhythms. Fig. 11 shows the distribution of average amplitudes of thalamic spindles as a function of the phase of the cortical slow oscillation. Spindle amplitudes are highest between slow oscillation phases 0 and *π*/4. The phase-amplitude relationship was indeed significant as shown by both tested measures (*KL–MI* = 0.0109, *p* < 0.001 and *MVL* = 11.3136, *p* < 0.001), while the phase-phase relationship cannot be deemed significant (*PLV* = 0.0020, *p* = 0.372 and *MI* = 0.0049, *p* = 0.266). We conclude that the thalamocortical model possesses significant phase-amplitude coupling between the phase of slow oscillations and the amplitude of thalamic spindles, but no phase-phase coupling was found.

**Figure 11:**
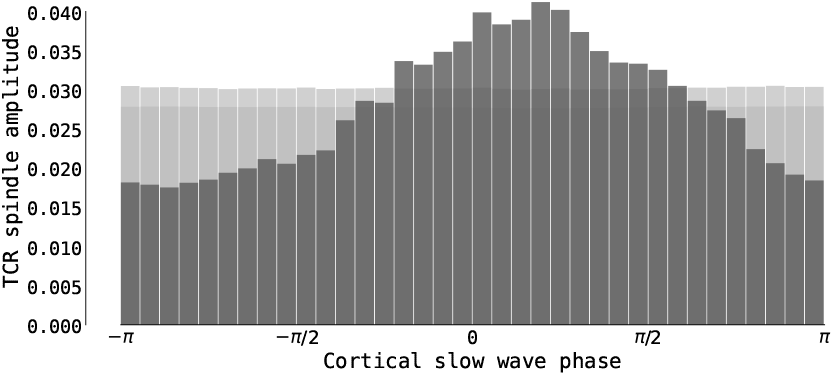
Phase-amplitude CFC in the thalamocortical motif. The figure show the distribution of the average amplitudes of the thalamic spindle dependent on the phase of the cortical slow oscillations in the simulation with short cortical UP states (time series from Fig. 10 with *μ_E_* = 0.61 nA). Dark grey bars denote the distribution in the simulated thalamocortical model, while light grey bars denote the mean and 95^th^ percentile distribution computed using 1000 IAAFT surrogates (for details, see Cross-frequency coupling (CFC) measures).

## 4. Discussion

This study investigated the dynamical states of a biophysically realistic neural mass model of a thalamocortical motif. To assess the contribution of each part of the model to the dynamics, we started with an isolated cortical and thalamic node and examined their individual dynamical landscapes. We perturbed each node with an external stimulus resembling their counterpart, i.e., the cortical model was perturbed by stimulation with spindle-like oscillations, and the thalamic model was perturbed using a square pulse with a low frequency, mimicking the idealized UP and DOWN state sequence of cortical SO activity. Next, we connected the isolated nodes to a full thalamocortical network and focused on spindle imprinting onto the cortical UP state activity and the interactions between spindles and slow oscillations, both being hallmark activity in the human brain during slow-wave sleep.

The results of our modeling study are in line with previous neuroimaging studies focusing on various aspects of sleep spindles and their interactions with cortical slow oscillations. In particular, the frequency of the spindles in our model (see Figs. 10A/C and C.3A/C) both in thalamic and cortical nodes matches the experimental values for fast spindles with an average of around 12 Hz [81, 75, 82, 83]. Typical spindle duration between 0.5–1 second also agrees with experimental values [75]. Finally, recent neuroimaging studies showed that sleep spindles possess a significant phase-amplitude relationship with cortical slow oscillations. More concretely, waxing periods of spindles are nested in the cortical UP states, and the spindle peak typically occurs right at (or just after) the slow oscillation UP state peak [41, 42, 84]. The presented thalamocortical model well reproduces this SO–spindle relationship (see Fig. 10B/D circular histograms and Fig. 11).

By dissecting the thalamocortical model, we sought the mechanistic explanation of the observed SO–spindle relationship. From the modeling perspective, we needed to recognize the difference between two different cortical parametrizations and their mechanistic consequences. The causal direction follows the path cortex → thalamus in the cortical parametrization with predominantly UP state activity with irregular, noise-driven DOWN state excursions. A sustained UP state in the cortex suppresses spindling via the corticothalamic connections since the TCR population cannot reach the necessary hyperpolarization (cf. Fig. 2). On irregular cortical DOWN swings caused by the background noise, the cortex is pushed into the DOWN state, which creates a window of opportunity for the thalamic DOWN state and a subsequent spindle waxing period. Finally, this spindle is then imprinted onto the cortical activity due to the thalamocortical projections. Reversely, with DOWN state-dominant cortical activity with irregular UP state excursions (cf. Fig. 4B middle), the causal direction reverses. In this case, without sustained cortical drive, the thalamus can generate “free” spindles, which subsequently push cortical activity into the UP state.

We left some parameters untouched despite exploring the model dynamics with various parameter settings. In particular, the conductances of calcium T-type currents *g_T_* in both thalamic populations control the underlying frequency of spindles in the spindle band (cf. Fig. 2). Allowing heterogeneous connections from the excitatory population in the cortical node to TCR and TRN populations (see Fig. 1) would control the balance between cortically driven excitation and inhibition, leading to an altered shape of the thalamic spindles. In particular, for a higher TRN/TCR input ratio, i.e., higher strength for cortex → TRN connection, the waxing periods are short—individual spindles contain only one or two oscillations. The change in shape is visible for higher ratios than 10 (not shown), and thus we have not included this parameter in our overall investigation. Note that the underlying frequency in the spindle band is not modulated by heterogeneous connection strengths between cortex and thalamus. Finally, one of the caveats of our study is the unknown realistic connection strengths in a thalamocortical motif in humans, which is why we treated both connection strengths from cortex → thalamus and thalamus → cortex as free parameters of our investigation. Given the maximum firing rates of both populations (cortex ~40 Hz, thalamus ~400 Hz), we indeed expected the connectivity strengths in both directions to have approximately a tenfold difference.

The dynamics of slow oscillations and spindles vary significantly between distinct sleep stages [12, 7, 80]. Various neuromodulators, such as acetylcholine (ACh) or histamine (HA), are known to vary significantly during sleep and awake, as well as across sleep stages [85, 86]. Specific effects of these neuromodulators can be implemented by changing the strength of the intrinsic and synaptic currents in the cortical and thalamic populations. In their biophysically realistic thalamocortical model, Krishnan et al. [6] identified a minimal set sufficient to account for characteristic changes of the brain’s electrical activity across the sleep-wake cycle. We have not included these effects and focused solely on the N3 sleep stage. Briefly, a reduction of ACh can be implemented by an increase of potassium leak conductance (*g_LK_*) in the thalamic node and an increase in spike-frequency adaptation (*b*) in the cortical node [87], while the effect of HA can be implemented as a shift in the activation curve of a hyperpolarization-activated current *I_h_* in the TCR [88].

This study merged two mass modeling approaches: a mean-field approximation of the Fokker-Planck equations for the cortical dynamics [43] and a mass model based on conductance-based average membrane voltage dynamics for the thalamus [34], and as such, showcases the applicability of hybrid modeling approaches. We are not aware of any caveats of this approach, as long as a natural link in the sense of coupling variable between the frameworks can be established. In our case, both frameworks operate with the notion of firing rate and use firing rate as the main model output and a coupling variable between their subpopulations. Using the already probed cortical node facilitates studying biophysically realistic stimulation protocols of the thalamocortical model in our future work. Our motivation was twofold: the cortical model introduced by Cakan et al. [43], which we used here, is biophysically realistic and very well studied with respect to its dynamical states. Moreover, a previous study also probed biophysically realistic stimulation protocols on par with transcranial direct current stimulation (tDCS) during sleep [43]. Finally, by merging different mass modeling approaches in our current work, we are also setting a stage for future extensibility of the thalamocortical model by including more cortical nodes, or, alternatively, nodes simulating different brain areas, e.g., the hippocampus.

One of the extension possibilities lies in topographic mass models. A well-known property of cortical spindles is their heterogeneity, i.e., mainly the difference between faster and slower spindles, where faster spindles are usually observed in parietal regions, while slower spindles are found in the frontal regions [89, 80, 9]. The main reason for this heterogeneity is the core and matrix thalamocortical pathways [90, 91]: core thalamocortical neurons are spatially selective and topographically organized, target a single cortical area, and project mainly to the granular layer. On the other hand, matrix neurons have diffuse, multiarea projections, characterized by multiple distant arbors, and reach mostly superficial layers of the cortex [90, 91]. The advantage of mass models over spiking models is the computational efficiency and the fast transition into layer-resolving 2D models by modeling more cortical layers with the same base model and a different between-layer corticocortical and corticothalamic connectivity and fan-outs [92], accounting for distinct core and matrix thalamocortical projections.

As already mentioned above, a realistic outlook and extension possibility is taking a step further in the direction of whole-brain dynamics and creating a whole-brain model consisting of many cortical nodes coupled using the structural connectome with the thalamus, able to undergo stimulation. Recently, Cakan et al. [76] constructed a deep sleep whole-brain model and studied the dynamics of local and global slow oscillation events. Each node in their model consisted of an excitatory-inhibitory pair of the mean-field approximation model of the AdEx neurons. Hence we might build on our current investigation of the thalamocortical motif.

We believe that our current investigation of the computationally efficient thalamocortical microcircuit model allows us to dive deeper into the sleeping brain and to shed light on the exact temporal structure and interaction of sleep rhythms involved in episodic memory consolidation.

## 5. Acknowledgments

Some of the computational resources used in this study were supplied by the project “e-Infrastruktura CZ” (e-INFRA LM2018140) supported by the Ministry of Education, Youth and Sports of the Czech Republic. N.J. was funded by the Operational Programme Research, Development and Education, Ministry of Education, Youth and Sport of the Czech Republic (co-funded by the EU) — Project Number CZ.02.2.69/0.0/0.0/19_074/0016209 (“*Modelling the sleeping brain: towards a neural mass model of sleep rhythms and their interactions*”) and by the Czech Science Foundation project number 21-32608S. This work was funded by the Deutsche Forschungsgemeinschaft (DFG, German Research Foundation) — Project number 327654276–SFB 1315 (C.C. and K.O.).

## 6. Conflict of interest

The authors declare that there is no conflict of interest.

## Appendix A. Precomputed transfer functions of the cortical model

**Figure A.1:**
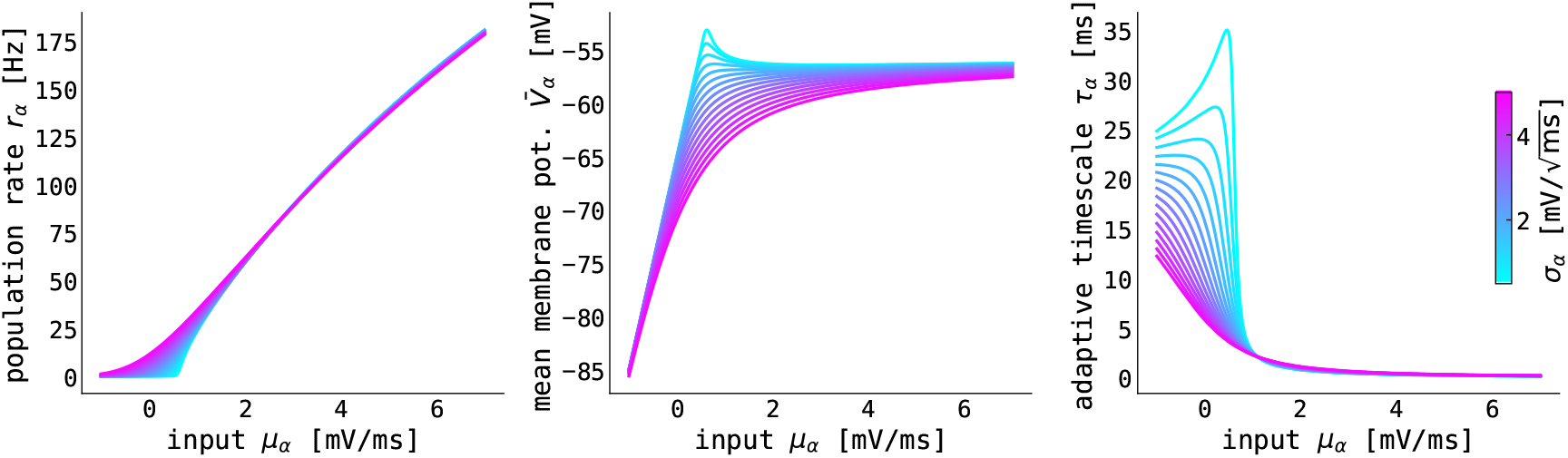
Precomputed quantities of the linear-nonlinear cascade model. Panels show (left to right) the nonlinear transfer functions Φ_*r*_ for the population firing rate *r_α_* (cf. Eq. 16), Φ_*V*_ for the mean membrane voltage 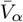 (cf. Eq. 17), and Φ_*τ*_ for the adaptive timescale *τ_α_* (cf. Eq. 18). The color coding represents the level of input current variance *σ_α_* across the population. The linear-nonlinear cascade was precomputed with the following single AdEx neuron parameters (see Eqs. (11) and (12)): membrane capacitance *C_m,c_* = 200 pF, leak conductance *g_L_* = 10 nS, membrane time constant *τ_m_* = *C*/*g_L_* = 20 ms, leak reversal potential *E_L_* = −65 mV, threshold slope factor Δ_*T*_ = 1.5 mV, spike initiation voltage threshold *V_T_* = −50 mV, spike voltage threshold *V_s_* = −40 mV, reset voltage *V_r_* = −70 mV, refractory period *T_ref_* = 1.5 ms. Note, that the somatic adaptation current (Eq. (13)) is not included in the linear-nonlinear cascade precomputation.

## Appendix B. Cortical model with finite noise driven by thalamic input

**Figure B.2:**
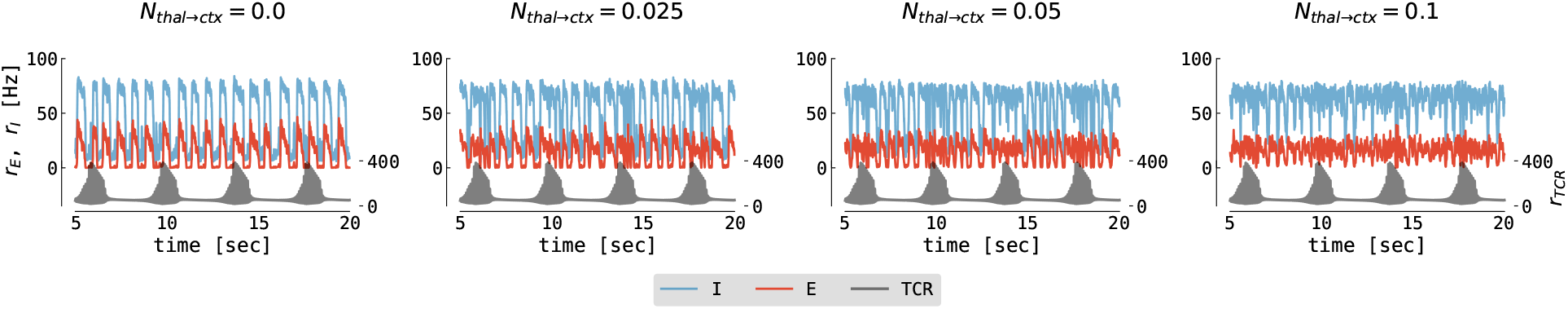
Cortical response to thalamic spindle stimulation with cortical noise. The panels show the firing rates of the excitatory cortical population (red), the inhibitory cortical population (blue), and the thalamic input (black) as function of time. The baseline in DOWN states for excitatory (inhibitory) population is around 0 Hz (50 Hz). Cortical model is parametrized at the border of the limit cycle and the UP state (*μ_E_* = 0.65 nA). Different columns show distinct values of thalamus → cortex connection strength ranging from *N_thal→ctx_* = 0.0 to *N_thal→ctx_* = 0.1. The parameters for the cortical node were: *μ_I_* = 0.4 nA, *σ_E_* = *σ_I_* = 0.05 mV/ms^3/2^, for all other parameters see Table G.2. The parameters for thalamic node were: *g_LK_* = 0.031 mS/cm^2^, *g_h_* = 0.062 mS/cm^2^, *σ_TCR_* = 0.0 mV/ms^3/2^, for all other parameters see Table G.1. The parameters of connected thalamocortical model are given in Table G.3.

## Appendix C. Slow oscillation–spindle interaction in the UP state regime with long UP states

**Figure C.3:**
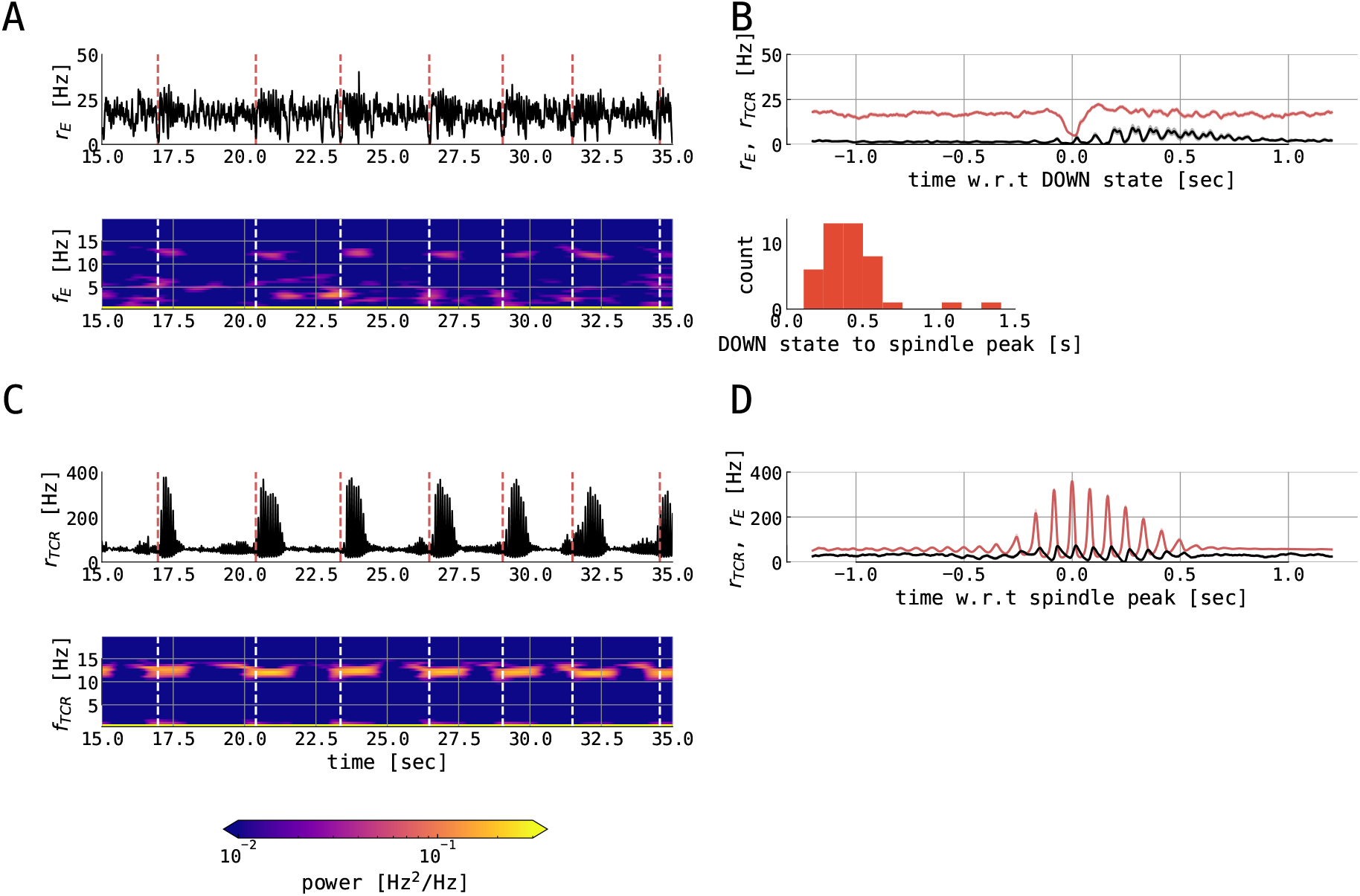
Spindles in the thalamocortical motif with long UP states. The figure shows various variables from 120 second of simulation of full thalamocortical model simulation with the cortical node being parametrized at the right border between the slow limit cycle and the UP state, further away from the bifurcation line (*μ_E_* = 0.66 nA). Individual panels show: (**A**) 20 seconds time series excerpt of the firing rate of the excitatory population in the cortical node (black) and its slow oscillation phase (blue) computed using the Hilbert transform of low-pass filtered cortical excitatory firing rate. The panel below shows the time-frequency representation of the cortical excitatory firing rates computed using the Short Time Fourier Transform with a 2 second time window. Dashed vertical lines (red in time series plot and white in time-frequency plot) denote the midpoints of cortical DOWN states which are followed by a cortical spindle in the 1.5 second window. (**B**) the mean ± the SEM of the cortical excitatory firing rates (red) and TCR firing rates (black), locked on the cortical DOWN states for the whole interval of 120 seconds. (**C**) 20 seconds time series excerpt of the firing rate of the thalamocortical relay population in the thalamic node (black). The panel below shows the time-frequency representation of the TCR firing rates computed using the Short Time Fourier Transform with a 2 second time window. Dashed vertical lines (red in time series plot and white in time-frequency plot) denote the midpoints of cortical DOWN states which are followed by a cortical spindle in the 1.5 second window. (**D**) the mean ± the SEM of the TCR firing rates (red) and cortical excitatory firing rates (black), locked on the thalamic spindle peaks in the whole interval of 120 seconds. The panel below shows the distribution of delays between midpoints of cortical DOWN states and spindle-band peaks. Note, that the the long UP states hinder slow oscillation phase estimation, since in a particularly long UP state, the Hilbert phase resets and no longer maps onto DOWN states with *ϕ_ctx,SO_* = −*π* = *π* and UP states with *ϕ_ctx,SO_* = 0. For this reason we do not quantify the cortical slow oscillation phase. The model was simulated with *N_thal→ctx_* = 0.12, *N_ctx→thal_* = 1.2, *μ_I_* = 0.4 nA, *σ_E_* = *σ_I_* = 0.05 mV/ms^3/2^, *σ_TCR_* = 0.005 mV/ms^3/2^, while other parameters were kept constant as per tables G.1, G.2, and G.3.

## Appendix D. Slow oscillation–spindle interaction in the DOWN state regime

**Figure D.4:**
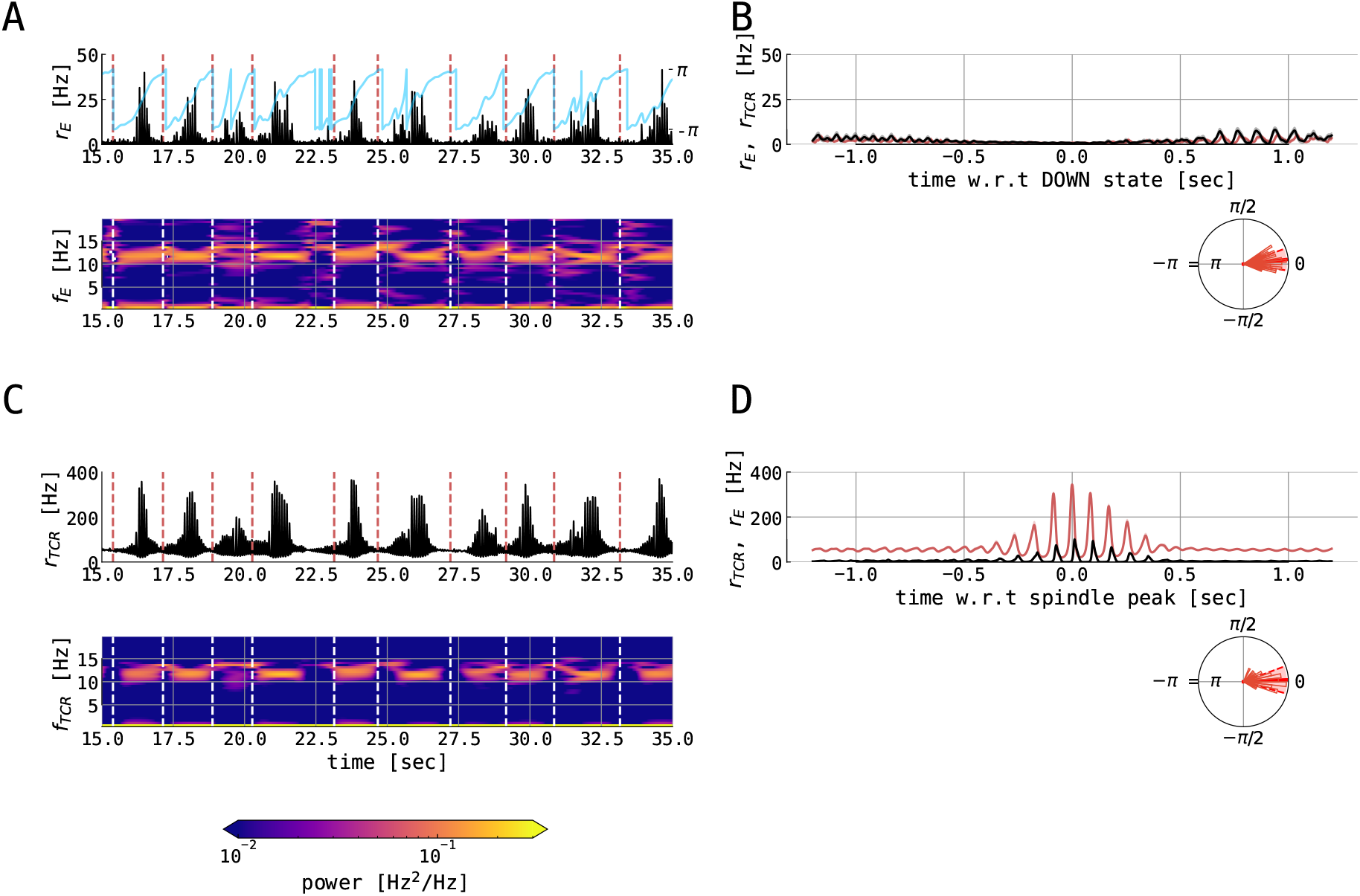
Spindles in the thalamocortical motif with the DOWN state parametrization. The figure shows various variables from 120 second of simulation of full thalamocortical model simulation with the cortical node being parametrized at the left border between the slow limit cycle and the DOWN state (*μ_E_* = 0.36 nA). Individual panels show: (**A**) 20 seconds time series excerpt of the firing rate of the excitatory population in the cortical node (black) and its slow oscillation phase (blue) computed using the Hilbert transform of low-pass filtered cortical excitatory firing rate. The panel below shows the time-frequency representation of the cortical excitatory firing rates computed using the Short Time Fourier Transform with a 2 second time window. Dashed vertical lines (red in time series plot and white in time-frequency plot) denote the midpoints of cortical DOWN states which are followed by a cortical spindle in the 1.5 second window. (**B**) the mean ± the SEM of the cortical excitatory firing rates (red) and TCR firing rates (black), locked on the cortical DOWN states for the whole interval of 120 seconds. (**C**) 20 seconds time series excerpt of the firing rate of the thalamocortical relay population in the thalamic node (black). The panel below shows the time-frequency representation of the TCR firing rates computed using the Short Time Fourier Transform with a 2 second time window. Dashed vertical lines (red in time series plot and white in time-frequency plot) denote the midpoints of cortical DOWN states which are followed by a cortical spindle in the 1.5 second window. (**D**) the mean ± the SEM of the TCR firing rates (red) and cortical excitatory firing rates (black), locked on the thalamic spindle peaks in the whole interval of 120 seconds. The panel below shows the distribution of cortical slow oscillation phases for the maximum of the thalamic *σ*-band peak. Shown are histogram bars in thin red and circular mean (± circular STD) in thick (dashed) red. The model was simulated with *N_thal→ctx_* = 0.12, *N_ctx→thal_* = 1.2, *μ_I_* = 0.4 nA, *σ_E_* = *σ_I_* = 0.05 mV/ms^3/2^, *σ_TCR_* = 0.005 mV/ms^3/2^, while other parameters were kept constant as per tables G.1, G.2, and G.3.

## Appendix E. Slow oscillation–spindle interaction in the limit cycle regime

**Figure E.5:**
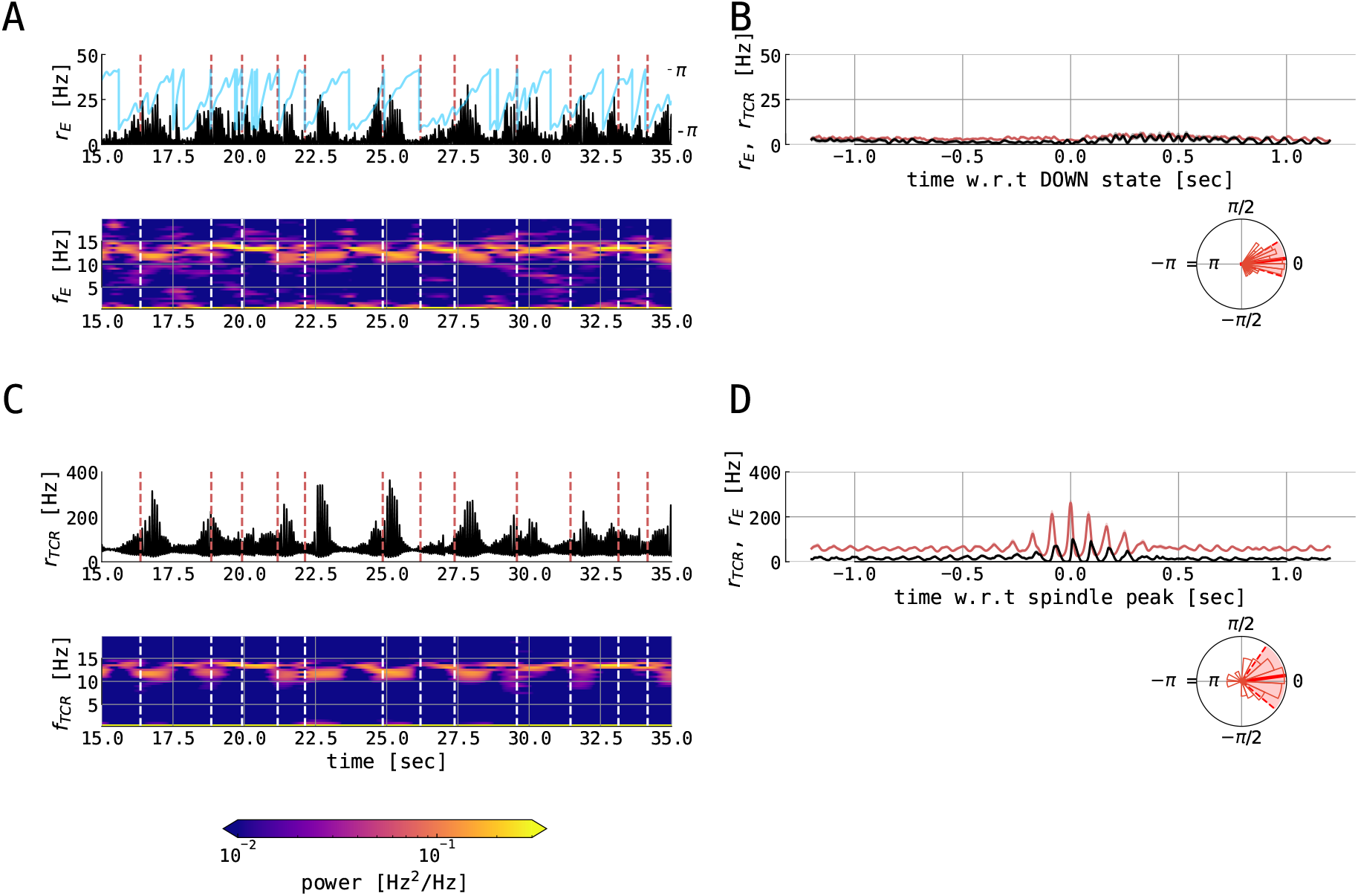
Spindles in the thalamocortical motif with the limit cycle parametrization. The figure shows various variables from 120 second of simulation of full thalamocortical model simulation with the cortical node being parametrized inside the slow limit cycle (*μ_E_* = 0.42 nA). Individual panels show: (**A**) 20 seconds time series excerpt of the firing rate of the excitatory population in the cortical node (black) and its slow oscillation phase (blue) computed using the Hilbert transform of low-pass filtered cortical excitatory firing rate. The panel below shows the time-frequency representation of the cortical excitatory firing rates computed using the Short Time Fourier Transform with a 2 second time window. Dashed vertical lines (red in time series plot and white in time-frequency plot) denote the midpoints of cortical DOWN states which are followed by a cortical spindle in the 1.5 second window. (**B**) the mean ± the SEM of the cortical excitatory firing rates (red) and TCR firing rates (black), locked on the cortical DOWN states for the whole interval of 120 seconds. (**C**) 20 seconds time series excerpt of the firing rate of the thalamocortical relay population in the thalamic node (black). The panel below shows the time-frequency representation of the TCR firing rates computed using the Short Time Fourier Transform with a 2 second time window. Dashed vertical lines (red in time series plot and white in time-frequency plot) denote the midpoints of cortical DOWN states which are followed by a cortical spindle in the 1.5 second window. (**D**) the mean ± the SEM of the TCR firing rates (red) and cortical excitatory firing rates (black), locked on the thalamic spindle peaks in the whole interval of 120 seconds. The panel below shows the distribution of cortical slow oscillation phases for the maximum of the thalamic *σ*-band peak. Shown are histogram bars in thin red and circular mean (± circular STD) in thick (dashed) red. The model was simulated with *N_thal→ctx_* = 0.12, *N_ctx→thal_* = 1.2, *μ_I_* = 0.4 nA, *σ_E_* = *σ_I_* = 0.05 mV/ms^3/2^, *σ_TCR_* = 0.005 mV/ms^3/2^, while other parameters were kept constant as per tables G.1, G.2, and G.3.

## Appendix F. Full set of model equations for the thalamic module

The complete mathematical description of the thalamic node in our model was taken from Costa et al. [34] and reads:

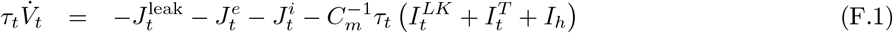

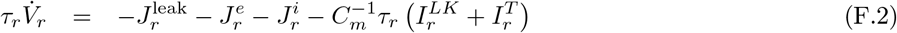

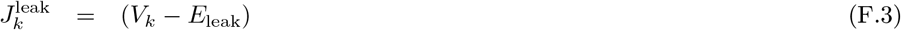

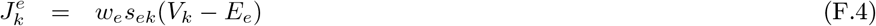

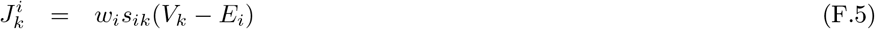

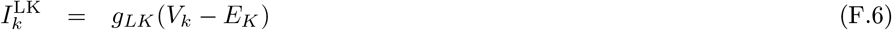

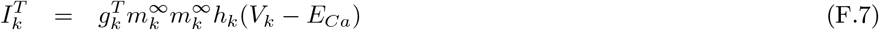

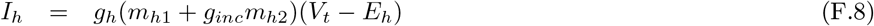

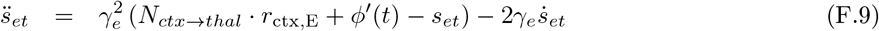

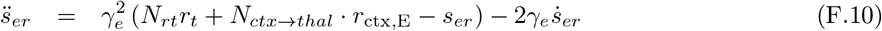

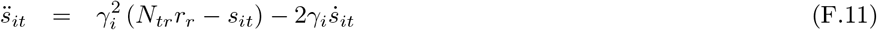

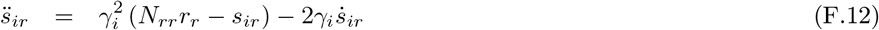

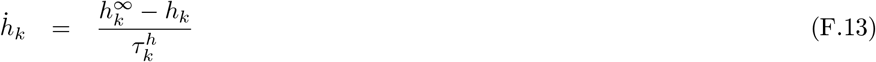

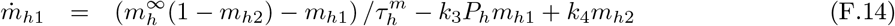

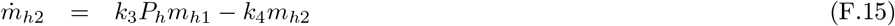

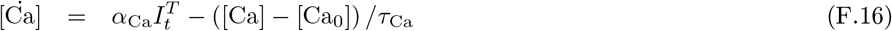

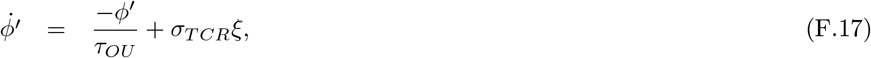

with subscripts *k* ∈ {*r, t*} with *r* standing for TRN and *t* for TCR population. Subscript *e* (*i*) denotes excitatory (inhibitory) synaptic type. *ξ* is drawn from random Gaussian white noise process with zero mean and unit variance. The gating functions are given by

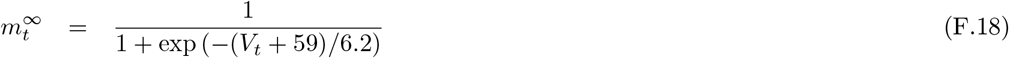

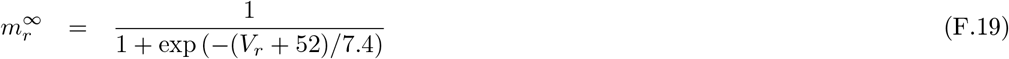

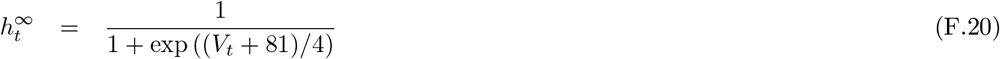

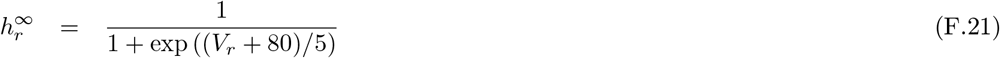

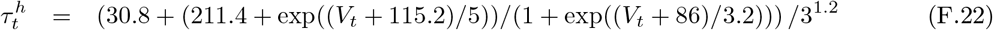

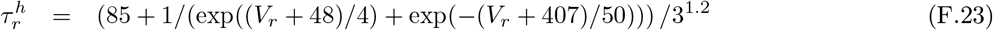

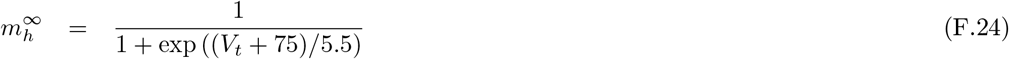

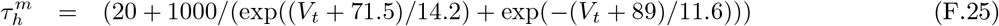

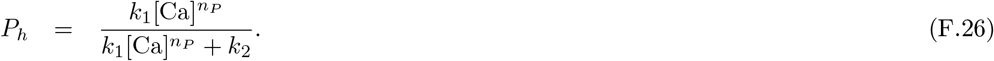

And finally, the firing rate transfer function obeys

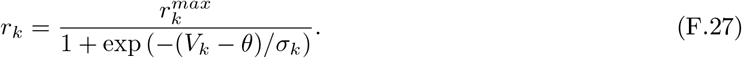

## Appendix G. Thalamocortical model parameters

**Table G.1:** Thalamic node parameters. Taken from Costa et al. [34].

| Parameter | Value | Description |
| --- | --- | --- |
| $C_m$ | $1 \mu\text{F}/\text{cm}^2$ | membrane capacitance |
| $\tau_t, \tau_r$ | 20 ms | membrane time constant of TCR and TRN |
| $r_t^{\max}, r_r^{\max}$ | 0.4 kHz | maximal firing rate of TCR and TRN |
| $\theta$ | -58.5 mV | firing threshold (half activation) |
| $\sigma_t, \sigma_r$ | 6 mV | inverse neural gain |
| $\gamma_e$ | 0.07 kHz | synaptic rate constant of AMPA synapses |
| $\gamma_i$ | 0.1 kHz | synaptic rate constant of GABA synapses |
| $N_{tr}$ | 5 | connectivity constant TRN → TCR |
| $N_{rt}$ | 3 | connectivity constant TCR → TRN |
| $N_{rr}$ | 25 | recurrent connectivity constant in TRN |
| $w_e, w_i$ | 1 ms | synaptic weights |
| $g_t^T$ | $3 \text{ mS}/\text{cm}^2$ | conductance of Ca current in TCR |
| $g_r^T$ | $2.3 \text{ mS}/\text{cm}^2$ | conductance of Ca current in TRN |
| $g_h$ | $[0.0-0.08] \text{ mS}/\text{cm}^2$ | conductance of rectifying current in TCR |
| $g_{LK}$ | $[0.0-0.08] \text{ mS}/\text{cm}^2$ | conductance of K leak current in TCR and TRN |
| $E_{\text{leak}}$ | -70 mV | Nernst reversal potential for leak channels |
| $E_e$ | 0 mV | Nernst reversal potential of AMPA channels |
| $E_i$ | -70 mV | Nernst reversal potential of GABA channels |
| $E_{Ca}$ | 120 mV | Nernst reversal potential of Ca channels |
| $E_K$ | -100 mV | Nernst reversal potential of K channels |
| $E_h$ | -40 mV | Nernst reversal potential of rectifying channels |
| $\alpha_{Ca}$ | $-51.8 \cdot 10^{-6} \text{ mM}/\text{mAms}$ | Ca influx rate |
| $\tau_{Ca}$ | 10 ms | Ca time constant |
| $[\text{Ca}_0]$ | $2.4 \cdot 10^{-4} \text{ mM}$ | Ca resting state concentration |
| $k_1$ | $2.5 \cdot 10^7 \text{ kHz}$ | } reaction velocities of rectifying current |
| $k_2$ | $4.0 \cdot 10^{-4} \text{ kHz}$ | |
| $k_3$ | $1.0 \cdot 10^{-1} \text{ kHz}$ | |
| $k_4$ | $1.0 \cdot 10^{-3} \text{ kHz}$ | |
| $n_P$ | 4 | number of Ca binding sites |
| $g_{inc}$ | 2.0 | conductivity scaling of rectifying current |

**Table G.2:** Cortical node parameters. Taken from Cakan et al. [43].

| Parameter | Value | Description |
| --- | --- | --- |
| $\sigma^{\text{ext}}$ | 1.5 mV/$\sqrt{\text{ms}}$ | standard deviation of external input |
| $K_E$ | 800 | number of excitatory inputs per neuron |
| $K_I$ | 200 | number of inhibitory inputs per neuron |
| $K_{E,\text{global}}$ | 250 | number of excitatory inputs from each global area |
| $c_{EE}, c_{IE}$ | 0.3 mV/ms | maximum AMPA PSC amplitude |
| $c_{II}, c_{EI}$ | 0.5 mV/ms | maximum GABA PSC amplitude |
| $c_{\text{global}}$ | 0.4 mV/ms | postsynaptic PSC amplitude for global connectome |
| $J_{EE}$ | 2.43 mV/ms | maximum synaptic current E→E |
| $J_{IE}$ | 2.6 mV/ms | maximum synaptic current E→I |
| $J_{EI}$ | -3.3 mV/ms | maximum synaptic current I→E |
| $J_{II}$ | -1.64 mV/ms | maximum synaptic current I→I |
| $\tau_{s,E}$ | 2 ms | excitatory synaptic time constant |
| $\tau_{s,I}$ | 5 ms | inhibitory synaptic time constant |
| $d_E$ | 4 ms | synaptic delay to excitatory neurons |
| $d_I$ | 2 ms | synaptic delay to inhibitory neurons |
| $\tau_m$ | $C/g_L = 20$ ms | membrane time constant |
| $a$ | 0 nS | subthreshold adaptation conductance |
| $b$ | 15 pA | spike-triggered adaptation increment |
| $E_A$ | -80 mV | Nernst reversal potential for adaptation current |
| $\tau_A$ | 1000 ms | adaptation current time constant |

**Table G.3:** Connected model parameters. Almost all parameters here were subject to change during our investigation. This table contains the typical values of respective parameters. Different parameter values than the ones written here are mentioned in the text and the respective figures captions. In the case of parameters which change dynamical properties of the model, we indicate the range of possible values.

| Parameter | Typical value | Description |
| --- | --- | --- |
| $N_{ctx \rightarrow thal}$ | 1.2 | connectivity strength cortex → thalamus |
| $N_{thal \rightarrow ctx}$ | 0.12 | connectivity strength thalamus → cortex |
| $d_{ctx,thal}$ | 13 ms | synaptic thalamocortical delay |
| $\tau_{OU}$ | 5 ms | time-scale of the Ornstein–Uhlenbeck process |
| $\sigma_E$ | [0.0–0.05] mV/ms<sup>3/2</sup> | noise variance for the cortical E population |
| $\sigma_I$ | [0.0–0.05] mV/ms<sup>3/2</sup> | noise variance for the cortical I population |
| $\sigma_{TCR}$ | [0.0–0.005] mV/ms<sup>3/2</sup> | noise variance for the thalamic TCR population |
| $\mu_E$ | [0.0–5.0] mV/ms | mean drift of the Ornstein–Uhlenbeck process for cortical E population |
| $\mu_I$ | [0.0–5.0] mV/ms | mean drift of the Ornstein–Uhlenbeck process for cortical I population |
| $\mu_{TCR}$ | 0 mV/ms | mean drift of the Ornstein–Uhlenbeck process for thalamic TCR population |
| $\xi$ | $\sim \mathcal{N}(0, 1)$ | Gaussian white noise process |

## Notes

### Competing Interest Statement

The authors have declared no competing interest.

